# Continental diversification and insular speciation in a widespread passerine (*Troglodytes musculus*) in southern South America

**DOI:** 10.1101/2025.11.07.687267

**Authors:** Maria Recuerda, Cecilia Kopuchian, Pablo Andrés Fracas, Pablo Luis Tubaro, Leonardo Campagana, Darío Alejandro Lijtmaer

## Abstract

Understanding how the evolutionary dynamics of geographically widespread species play out in space and time is critical for uncovering the processes shaping biodiversity. We integrated mitochondrial DNA and genome-wide SNPs to investigate the history of diversification of the Southern House Wren (*Troglodytes musculus*) in southern South America and its insular relative, Cobb’s Wren (*T. cobbi*), from the Malvinas/Falkland Islands (MFI). *T. musculus* is an abundant and widespread species, and by studying its population structure and demographic history we aim to uncover the processes that may have shaped the evolutionary history of this and other co-distributed taxa in southern South America. All analyses indicate a latitudinal divergence, with an initial split between northern populations (Bolivia, Northern and Central Argentina, Uruguay) and southern populations from Patagonia and the MFI. Shortly after this initial divergence, further diversifications occurred, including the colonization of the MFI from Patagonia, where this newly formed insular population became isolated from the continent, and the split between an Andean lineage and a widespread, lowland lineage occurring in lowland Bolivia, Northern and Central Argentina and Uruguay. Both the timing of these splits and the mitochondrial and genomic patterns of variation uncovered in *T. musculus* are consistent with a major role of glaciations -and particularly the Great Patagonian Glaciation-in the diversification of this species in the southern Neotropics. The negligible gene flow between the MFI and the continent, which derived in the formation of a new species, Cobb’s Wren, contrasts with the notorious gene flow in the secondary, post-glacial contact zones between the northern and southern lineages, although with reduced gene flow between the Andean and lowland populations. We also observed cases of mito-nuclear discordance, highlighting the value of multi-locus approaches in revealing complex evolutionary histories. These results underscore how historical divergence, ecological barriers and corridors, and secondary contact after varying degrees of differentiation in allopatry jointly shape population structure and genetic diversity, with implications for understanding the diversification of widely distributed taxa.

## Introduction

Phylogeographic studies have revealed that while many species exhibit genetic subdivisions coincident with geographic barriers, patterns of genetic structure are often also shaped by less evident ecological traits such as migratory behavior, habitat preferences, and historical demographic shifts (Emerson and Hewitt 2005). Such patterns suggest that evolutionary histories are often more complex than expected, particularly in highly mobile taxa (such as birds) with broad distributions. Understanding the extent and nature of population structure is critical, as it bears on questions of taxonomy, speciation, and conservation (Coates et al. 2018). Species with wide geographic ranges can be leveraged to investigate how geography, past and current climate, and ecological factors collectively drive the diversification of both the focal taxon and other co-distributed species in the region, as well as how this might eventually lead to speciation. The advent of genomic data has transformed our understanding of these processes, revealing that gene flow during divergence is often more extensive than previously thought, and that natural and sexual selection are crucial forces in maintaining species boundaries despite ongoing genetic exchange (Feder et al. 2012; Martin et al. 2013; Seehausen et al. 2014).

In the Neotropics, several geographic barriers have been postulated as drivers of allopatric speciation in diverse taxa: the Andes mountains (Weir 2006; Elias et al. 2009; Lagomarsino et al. 2016; De la Riva et al. 2018), Amazonian rivers (Boubli et al. 2015; Pirani et al. 2019; Ribas et al. 2025) and unsuitable areas separating refuges during Pleistocene glacial periods (Sersic et al. 2011; Femenias et al. 2020; Sosa-Pivatto et al. 2020), among others. In particular, these have also been considered the main drivers of avian diversification. The uplift of the Andes provided a suitable habitat for the specialization of highland birds and generated a barrier for some lowland species, increasing avian diversification from a combination of vicariant and dispersal events (Brumfield and Edwards 2007; Weir and Price 2011; Hazzi et al. 2018). Similarly, major Amazonian rivers have long been recognized as barriers delimiting avian species ranges and areas of endemism (Sick 1967; Capparella 1988, 1991; Hayes and Sewlal 2004), with more recent studies showing genetic differentiation on opposite margins (Ribas et al. 2012; Naka and Brumfield 2018). Finally, glacial cycles have boosted speciation in southern South America and the Andes due to both the advance of ice sheets during glaciations and the repeated movement and fragmentation of suitable habitats during glacial cycles (Weir 2006; Jetz et al. 2012; Hazzi et al. 2018), but their role as a speciation driver at lower latitudes is more contentious (Cabanne et al. 2016; Thom et al. 2020; Guayasamin et al. 2024).

These geographic barriers have been traditionally considered to produce complete isolation among diverging populations, consistent with speciation in allopatry. However, it is now clear that this is not necessarily the case and that barriers can be permeable, both in the initial process of divergence between populations or afterwards, allowing for secondary contact and gene flow between the previously isolated lineages (Morales-Rozo et al. 2017; Acosta et al. 2021; Céspedes-Arias et al. 2021; Bukowski et al. 2024; Moncrieff et al. 2024; Weir et al. 2024). Moreover, speciation without the need for long periods of allopatry has also been documented for Neotropical birds, including cases of hybrid speciation (Barrera-Guzmán et al. 2018; Lamichhaney et al. 2018; Turbek et al. 2021). However, we still do not know how common these modes of speciation are, as well as the level of permeability of these reproductive barriers, and the factors leading to different outcomes when there is gene flow during -or after-the diversification process remain poorly understood (Weir et al. 2015, 2024; Cadena et al. 2016). In this regard, recent studies of Neotropical barriers that are common to multiple avian taxa show that the permeability of barriers is idiosyncratic and depends on the biological attributes of each species (Smith et al. 2014; Naka and Brumfield 2018; Kopuchian et al. 2020; van Els et al. 2021; Luszczak et al. 2025). Addressing these gaps is essential to fully comprehend the evolutionary processes that lead to speciation in the Neotropics.

The House Wren species complex (*Troglodytes aedon, T. musculus* and associated insular forms) exemplifies the challenges of understanding diversification in widespread taxa. It was traditionally considered a single species (*T. aedon*) and the passerine with the broadest distribution in the New World, ranging from southern Canada to Tierra del Fuego in southern South America and occupying a wide variety of habitats and elevations (Johnson 2024). However, recent taxonomic revisions have revealed substantial cryptic diversity within this complex, resulting in its split into nine distinct species based on morphological traits and geography (Chesser et al. 2024). In the first place, *T. musculus*, the Southern House Wren, is now recognized as a separate species encompassing all mainland populations from south of the Isthmus of Tehuantepec in southern Mexico to Tierra del Fuego, while populations north of this divide remain within *T. aedon*, the Northern House Wren (Chesser et al. 2024; Anderson et al. 2025). Similarly, island populations from Dominica, St. Lucia, Cozumel, St. Vincent, Grenada, Socorro, and Clarión have each been elevated to full species status (*T. martinicensis*, *T. mesoleucus*, *T. beani*, *T. musicus*, *T. grenadensis*, *T. sissonii*, and *T. tanneri*, respectively) (Klicka et al. 2023; Chesser et al. 2024). Additionally, Cobb’s Wren (*Troglodytes cobbi*), which is restricted to the Malvinas/Falkland Islands (MFI) in the Southern Atlantic Ocean, is recognized as a separate species based on significant morphological, ecological, genetic, and vocal divergence (Campagna et al. 2012; Chesser et al. 2024). These islands are found ∼400 km off the coast of Argentine Patagonia and have acted as a glacial refugium during the Pleistocene, promoting differentiation and speciation in several other birds (Campagna et al. 2012, 2019; Kopuchian et al. 2016) and at least one mammal (Austin et al. 2013).

Initial studies based on mitochondrial DNA offered valuable insights into the evolutionary history of this species complex in the Neotropics, revealing the presence of multiple divergent mitochondrial lineages at different geographic scales (Kerr et al. 2009; Campagna et al. 2012; Galen et al. 2015). However, incorporating nuclear markers is crucial to assess the isolation between these initially identified lineages while considering historical and ongoing gene flow. Recent genomic studies have begun to overcome this limitation. Klicka et al. (2023) found deep phylogeographic structure within the complex, showing not only strong genome-wide differentiation in island populations such as *T. beani*, *T. sissonii*, and *T. tanneri*, but also among mainland populations. Notably, South American populations of *T. musculus* showed additional substructure, revealing distinct groups in Ecuador/northern Peru, central/south-central Peru, and east of the Andes. Both nuclear genomic DNA and mitochondrial sequences supported the presence of strong phylogeographic structure, but their patterns — and thus the relationships they suggest among groups — were discordant (Klicka et al. 2023).

While these findings highlight the broad-scale complexity of the evolutionary history in this group, finer-scale population structure in South America remains understudied, especially in the southern cone of the continent. Previous studies from this region based on mitochondrial DNA showed the presence of multiple lineages, with up to 5% divergence in their cytochrome c oxidase subunit I (COI) sequences, but neither their relationships nor their degree of isolation have been assessed so far (Kerr et al. 2009; Campagna et al. 2012). To explore this further, we gathered genomic and mitochondrial data from *T. musculus* across southern South America, and *T. cobbi* from the MFI, to study their evolutionary history in this region. We emphasized the analysis of the patterns of gene flow, both in the mainland and between *T. musculus* and *T. cobbi*, and the role played by the glacial cycles, as they have been a key diversification factor for other southern taxa (Sersic et al. 2011; Femenias et al. 2020; Sosa-Pivatto et al. 2020; Acosta et al. 2021). This allowed us to disentangle the relationship between the southern lineages of *T. musculus* and to understand how their patterns fit into the broader evolutionary context of this species complex.

## Materials and Methods

### Study system and sampling collection

Individuals of *T. musculus* have been sampled broadly across southern South America by the research group of the Ornithology Division at the Museo Argentino de Ciencias Naturales “Bernardino Rivadavia” (MACN) for the last 20 years, with geographic coverage spanning most of Argentina (including Patagonia) and localities in highland and lowland Bolivia and in Uruguay (Figure 1A, Table S1). Collected specimens were deposited at the National Ornithology Collection at MACN, and samples of their pectoral muscle were deposited at the National Ultrafrozen Tissue Collection (preserved at −75 °C). In the case of individuals that were bled and released, blood samples were deposited at the National Ultrafrozen Tissue Collection (preserved in 100% ethanol at −75 °C) and photographic e-vouchers were obtained whenever possible. A total of 111 *T. musculus* individuals were used for this study (Figure 1A), including both muscle and blood samples (a fraction of these samples had already been used in previous studies of primarily mtDNA of the species; (Kerr et al. 2009; Lijtmaer et al. 2011; Campagna et al. 2012). In addition, *T. cobbi* was represented in the dataset by 11 individuals sampled in the MFI (Campagna et al. 2012; See Figure 1A).

**Figure 1.**
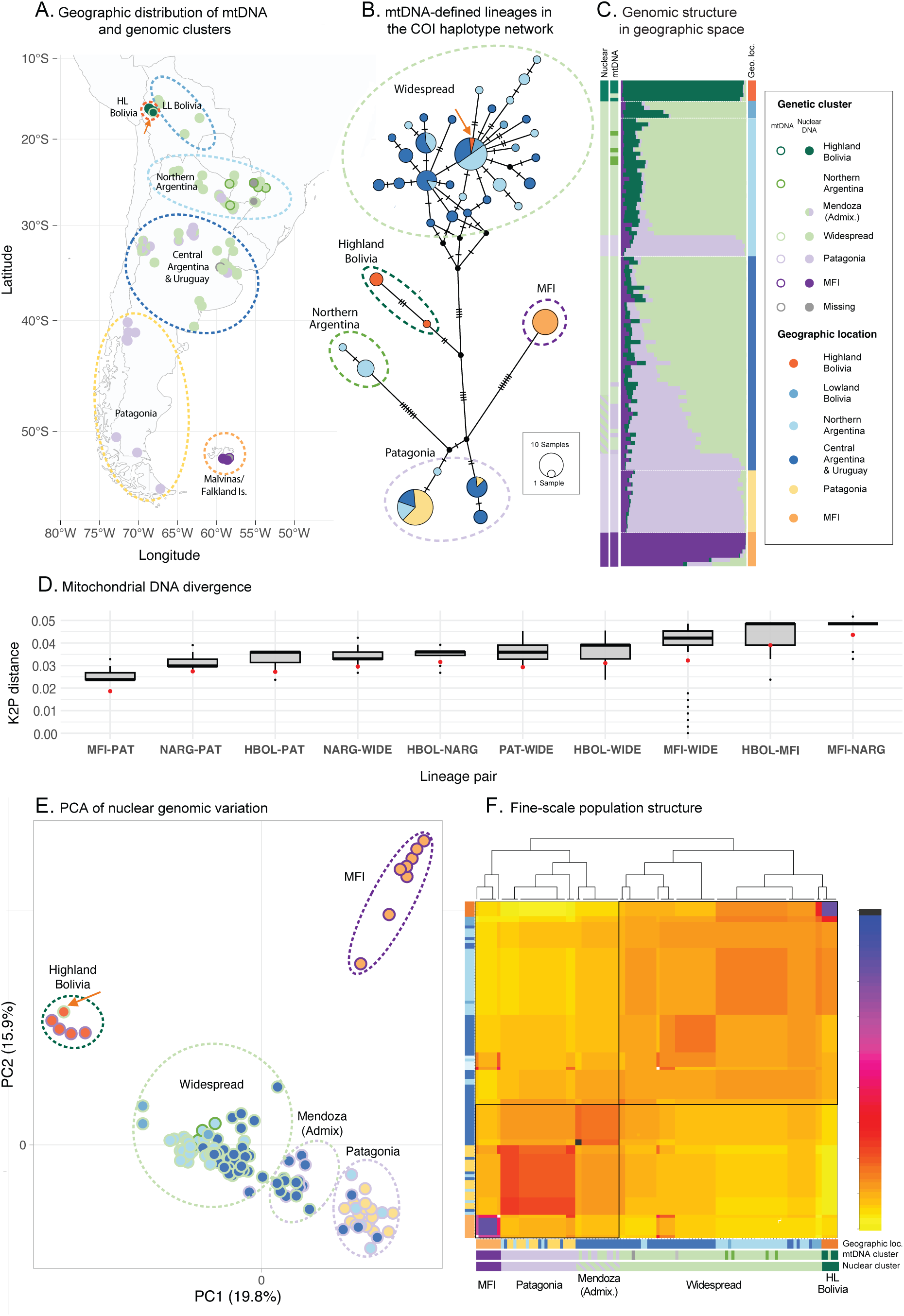
Geographic distribution and genetic structure of populations of the Southern House Wren across southern South America, and Cobb’s Wren in the Malvinas/Falkland Islands. **A)** Sampling locations colored by nuclear genetic cluster assignment (according to fineRADstructure, see panel E), with circle borders indicating mtDNA group (see panel B). Samples are grouped by major geographic regions: Highland Bolivia (HL Bolivia), Lowland Bolivia (LL Bolivia), Northern Argentina, Central Argentina & Uruguay, Patagonia, and the Malvinas/Falkland Islands. In localities where the dots overlap, they have been slightly displaced for visualization purposes. The orange arrow highlights an example of mito-nuclear discordance, as shown in panels A, B, and D: a Highland Bolivia individual carrying a Widespread mitochondrial DNA haplotype. **B)** Median-joining haplotype network of cytochrome oxidase subunit I (COI) sequences for 121 individuals from *Troglodytes musculus and T. cobbi*. Each circle represents a unique mitochondrial haplotype, with circle size proportional to the number of individuals sharing that haplotype (see scale inset). Circle colors indicate the geographic origin of individuals, as shown in the legend. Dashed ellipses group haplotypes by mtDNA genetic cluster. The length of the branches connecting haplotypes is proportional to the number of mutational steps between them, which are indicated by the number of line marks on each branch. **C)** STRUCTURE analysis showing ancestry proportions for individuals assuming four genetic clusters (K = 4), with individuals ordered by geographic location. The bar on the left shows the genetic clustering according to fineRADstructure. **D)** Boxplots showing the distribution of mitochondrial DNA (COI, Kimura 2-parameter) pairwise divergences between lineages. Each box represents the interquartile range (25th–75th percentiles), with the horizontal line indicating the median and whiskers extending to values within 1.5 × the interquartile range. Each lineage pair is plotted separately. The red dot indicates the net genetic divergence (D_a_ estimate) for each lineage pair. Population abbreviations: HBOL—Highland Bolivia; NARG—Northern Argentina; WIDE—Widespread; MFI—Malvinas/Falkland Islands; PAT—Patagonia. **E)** Principal component analysis (PCA) of genetic variation reveals distinct genetic clusters. Points are colored by geographic location, the borders are colored according to their mtDNA cluster, and dashed ellipses delineate nuclear genetic clusters. **F)** fineRADstructure analysis showing the pairwise co-ancestry matrix with hierarchical clustering (top) and population assignments by geography (left and bottom bar) and mtDNA and genomic cluster (bottom bars).

### DNA extraction, mitochondrial and ddRAD sequencing, and SNP genotyping

We used COI sequences as our mitochondrial DNA marker. Nineteen *T. musculus* sequences and the 11 *T. cobbi* sequences had been already obtained in previous studies (Kerr et al. 2009; Campagna et al. 2012). For the rest of the individuals, DNA extraction was performed following a silica-based protocol described by Ivanova et al. (2006) and the polymerase chain reaction (PCR) cocktail and thermocycling profile followed Lijtmaer et al. (2012). Primers used for the amplification of the 695 base pairs (bp) of the COI were BirdF1 (Hebert et al. 2004) and COIbirdR2 (Kerr et al. 2009). Sequencing was conducted either at the Centre for Biodiversity Genomics of the University of Guelph, Canada, or at Macrogen Korea and was performed bidirectionally with the same primers used for amplification. GenBank accession numbers for all COI sequences generated in this study are included in the Supporting information. Sequences were assembled and edited in Sequencher (Nishimura 2000) and Geneious Prime 2024.0.4 (https://www.geneious.com), obtaining a final trimmed alignment length of 695 bp.

To obtain genomic data, we extracted high-quality genomic DNA from the entire set of samples using the QIAGEN Blood and Tissue kit (Qiagen, Valencia, CA) following the manufacturer’s protocol. We prepared double-digest restriction site-associated DNA (ddRAD) libraries following the protocol adapted from Peterson et al. (2012), as detailed in Thrasher et al. (2018). Sequencing was performed together with samples from other projects on three Illumina HiSeq2500 (100 bp single end) lanes at the Cornell University Biotechnology Resource Center (BRC).

We used the FASTX-Toolkit (https://github.com/agordon/fastx_toolkit) to trim the 3′ end of all ddRAD reads to a length of 97 bp using FASTX Trimmer v.0.0.13. Additionally, we discarded reads if any base had a Phred score below 10, corresponding to 90% base call accuracy, or if more than 5% of bases had a Phred score below 20 (accuracy of 99.9%). We assessed the quality of raw and trimmed reads using FastQC v0.12.1(Andrews 2010) and then used MultiQC v1.15 (Ewels et al. 2016) to generate a comprehensive summary report. We aligned the trimmed reads to the Rufous Wren reference genome (*Cinnycerthia unirufa*, GCA_035048185.1_ASM3504818v1) using BWA-MEM2 v2.2.1(Vasimuddin et al. 2019). A genome for *Trolglodytes* is not publicly available, we therefore chose the reference genome based on completeness and phylogenetic proximity (Imfeld et al. 2024). After alignment, we converted the resulting SAM files to sorted BAM files, added read group information, and marked duplicate reads using Picard Tools v2.26.1 (Picard toolkit 2019). We then assessed alignment statistics using SAMtools v1.20 (Danecek et al. 2021) and evaluated mapping statistics with Qualimap v2.2.1 (Okonechnikov et al. 2016). Mapping rates averaged 84.63% for *T. musculus* and 90.81% for *T. cobbi*. For SNP calling and population genetic analyses, we used the Stacks v2.67 pipeline (Rochette et al. 2019) to assemble loci and identify SNPs from aligned reads. Using VCFtools v0.1.16 (Danecek et al. 2011), we applied filtering thresholds based on minor allele frequency, Hardy-Weinberg equilibrium (p < 0.0001), mean sequencing depth, and missing data. We retained only biallelic SNPs with a minor allele frequency above 0.01, a mean sequencing depth between 30 and 60, and a missing data threshold below 10%. Additionally, we removed individual ID9103 from Northern Argentina because after filtering it had more than 20% missing data. After mapping, we initially recovered 244,795 variable sites, which were reduced to 6,650 loci after quality filtering. The final dataset had an average sequencing depth and missing data proportion of 44.8x (±20.6 SD) and 2.6% ± (±0.03 SD), respectively.

After processing the 122 individuals of our dataset (111 *T. musculus*, 11 *T. cobbi*) and applying the quality criteria described above, we retained both ddRADseq and COI data from 115 individuals (107 *T. musculus*, 8 *T. cobbi*). For six individuals (3 *T. musculus* and 3 *T. cobbi*) we retained COI information and discarded their ddRADseq data due to high missingness. The opposite was true for one individual of *T. musculus* for which mitochondrial sequencing failed. The final dataset used for the analyses is included in Table S2.

### Haplotype network and mtDNA divergence

We constructed a network to visualize the relationships among the 121 COI mitochondrial haplotypes. The resulting alignment was imported into PopART v1.7 (Leigh et al. 2015), where a median-joining network was generated with the epsilon parameter set to zero to minimize complexity. This allowed us to explore haplotype sharing and divergence in a geographic framework.

We quantified mitochondrial DNA divergence by calculating pairwise genetic distances under the Kimura 2-parameter (K2P) model (Kimura 1980), implemented in the R package *ape* v5.7. Average distances were then computed to assess divergence levels. We also calculated net sequence divergence (D_a_; (Nei 1987) following Luszczak et al. (2025), which represents the mean genetic divergence between populations after accounting for within-population variation.

### Genome-wide population structure

We performed linkage disequilibrium (LD) pruning and principal component analysis (PCA) using PLINK2 (Chang et al. 2015). First, we applied LD-based SNP pruning with a sliding window of 50 SNPs, an incremental shift of 10 SNPs, and an r² threshold of 0.1, resulting in a final dataset of 3,820 unlinked variants. Next, we used the pruned SNP set to generate a binary PLINK file and conducted PCA to assess population structure.

We also analyzed population structure using STRUCTURE v2.3.4 (Pritchard et al. 2000) to assess genetic clustering and admixture among individuals. The VCF file of unlinked SNPs was converted to STRUCTURE format using PGDSPIDER v3.0.0.0 (Lischer and Excoffier 2012). Prior to running the main analyses, we inferred the lambda parameter and incorporated it into subsequent STRUCTURE runs. We performed five independent runs for each K value, ranging from one to seven, using the admixture model and correlated allele frequencies. Each run consisted of 250,000 burn-in iterations followed by 500,000 Markov Chain Monte Carlo (MCMC) iterations to ensure convergence. We did not use sampling locality as a prior (LOCDATA=0). To determine the most likely number of clusters (K), we used structureHarvester (Earl and VonHoldt 2012) to analyze results based on both the Evanno method (ΔK) (Evanno et al. 2005) and the mean log probability of the data [Mean LnP(Data)].

To investigate population relationships using haplotype-based methods, we used fineRADstructure v0.3 (Malinsky et al. 2018), which groups individuals based on shared haplotype segments. First, we calculated the coancestry matrix using RADpainter, which processed the haplotype data generated by STACKS v2.67. Next, we assigned individuals to populations with finestructure, running an MCMC simulation with 100,000 burn-in and the same number of sampling iterations, and a thinning interval of 1,000. A coancestry tree was then built using the tree-building mode of finestructure with 10,000 iterations. Finally, we visualized the results using the fineRADstructurePlot.R script in R, which generates heatmaps and dendrograms of the coancestry matrix.

### Isolation by distance

To test for isolation by distance (IBD), we conducted a Mantel test (Mantel 1967) comparing pairwise genetic and geographic distances among individuals. Genetic distances were calculated from genome-wide SNP data using a Euclidean distance matrix derived from a genlight object in the R package adegenet (Jombart 2008), based on genotypes extracted from a VCF file with vcfR (Knaus and Grünwald 2017). Geographic distances were calculated as pairwise great-circle distances (in kilometers) between sampling coordinates using the Haversine formula implemented in the geosphere package (Hijmans et al. 2024). The Mantel test was performed with the vegan package (Oksanen et al. 2013) using Pearson’s product-moment correlation and 999 permutations. To visualize the relationship, we generated a scatterplot of genetic versus geographic distances, with each point color-coded by its local two-dimensional density estimated via kernel density estimation (kde2d) to illustrate sampling density across the space of comparisons. To account for potential limitations of the Mantel test due to non-independence of pairwise distances, we also applied a Multiple Regression on Distance Matrices (MRM; (Guillot and Rousset 2013) using the ecodist package.

### Hybrid Index Analysis

We assessed whether the Mendoza population originated through admixture between two of the detected lineages, using the R package triangulaR (Omys-Omics, GitHub). Variant data were obtained from the filtered VCF file, retaining individuals belonging to the genetic clusters of Patagonia, Widespread, or of intermediate ancestry corresponding to the geographic locality of Mendoza. To identify informative sites, we applied an allele frequency difference filter between the Patagonia and Widespread populations. While a threshold of 0.9 is typically used to select highly differentiated sites, we had to lower this value to 0.5 to recover loci. We therefore tested two thresholds for the minimum allele frequency difference (0.5 and 0.4), which resulted in 5 and 11 sites passing each threshold, respectively. Due to the low number of markers obtained with the 0.5 threshold, which render this option uninformative, we kept the 0.4 threshold for the analysis. For each individual, we then calculated the hybrid index and interclass heterozygosity using the hybridIndex() function. Triangle plots were generated with the triangle.plot() function to visualize the distribution of individuals relative to expectations under different hybridization scenarios.

### Estimating effective migration and diversity surfaces (EEMS)

To identify potential barriers to gene flow, we used EEMS (Estimated Effective Migration Surfaces, v20181127; Petkova et al. 2016), which models migration using a stepping-stone framework where gene flow occurs only between neighboring demes arranged in a user-defined grid. Migration rates are estimated and interpolated across the demes to create a visual representation of genetic diversity in relation to geographic location. Regions where genetic similarity declines more rapidly than expected through isolation by distance alone indicate potential barriers to gene flow. We first ran runeems_snps at densities of 300-700 (in increments of 100) to determine an appropriate deme density. For each density, we conducted three independent runs with 5 million MCMC iterations, including 2 million burn-in iterations and applying a thinning of 9,999. The final diversity and migration surfaces were averaged across all runs and deme densities using rEEMSplots v0.0.2, following the recommendations of Petkova et al. (2016).

### Phylogenetic inference

We reconstructed maximum likelihood phylogenies from mitochondrial haplotypes and nuclear SNP data using both IQ-TREE v2.2.2.6 (Minh et al. 2020) and RAxML-NG v1.2.0 (Kozlov et al. 2019) to assess the robustness of the inferred topologies. For the mitochondrial analysis, in addition to the 121 COI sequences from our ingroup, we added two *Troglodytes aedon* sequences from Canada as outgroups, which we downloaded from GenBank (DQ434786.1 and DQ434787.1) and were originally generated by Kerr et al. (2007). Sequences were aligned in Geneious, exported in PHYLIP format, and analyzed with IQ-TREE, using ModelFinder to select the best-fit substitution model (-m MFP). Node support was estimated with 1,000 ultrafast bootstrap replicates (-bb 1000) and SH-aLRT tests (-alrt 1000). In parallel, we conducted phylogenetic inference in RAxML-NG under the GTR+G substitution model with 100 non-parametric bootstrap replicates.

For the nuclear dataset, we inferred relationships based on a panel of SNPs filtered for linkage disequilibrium. The reference genome was included as an additional individual to serve as the outgroup, and a multi-FASTA alignment was generated and converted to PHYLIP format in Geneious. We reconstructed the tree using IQ-TREE under the GTR+ASC model, with 1000 ultrafast bootstrap replicates (-B 1000) and the reference genome specified as the outgroup (-o REF). We additionally ran RAxML-NG using the GTR+GAMMA model with ascertainment bias correction, 100 bootstrap replicates, and 10 parsimony-based starting trees. The resulting topologies were visualized in FigTree v1.4.4 (http://tree.bio.ed.ac.uk/software/figtree/).

### Demographic inference using G-PhoCS

To infer historical demographic parameters, we used G-PhoCS v1.3 (Generalized Phylogenetic Coalescent Sampler; Gronau et al. 2011), a Bayesian framework that co-estimates population divergence times, effective population sizes, and migration rates from multiple loci. Input data were generated from a VCF file containing both variant and invariant sites, using the *populations* program in Stacks to export loci in FASTA format, filtered to include loci present in at least 90% of individuals. A custom Perl script was used to convert these FASTA alignments into the G-PhoCS input format, preserving two haplotypes per sample. We selected five individuals per population (ten haplotypes total) based on population assignments from STRUCTURE, leaving the admixed individuals from Mendoza out of the analysis. Although 7,929 loci passed filtering, a random subset of 5,000 loci was used for the final G-PhoCS analysis to reduce computational burden.

We fit an Isolation with Migration (IM) model for all pairwise population comparisons, allowing for estimates of effective population sizes, divergence times, and bidirectional migration rates between each population pair. Priors for population sizes, divergence times, and migration rates were drawn from gamma distributions with parameters tuned to ensure proper mixing of the MCMC chains. We ran one MCMC chain of 2,000,000 iterations, discarding the first 200,000 as burn-in and sampling every 100 iterations. Convergence was assessed visually and through Effective Sample Size (ESS) calculations using the coda package in R (Plummer et al. 2015). Demographic parameter estimates were summarized from the posterior distributions, and 95% highest posterior density (HPD) intervals were calculated. Population divergence times were scaled using a generation time of 1 year (Johnson 1998) and two available avian mutation rates to capture uncertainty in the estimates: 4.6 × 10⁻⁹ substitutions per site per generation, estimated for flycatchers (Smeds et al. 2016) and 1.2 × 10⁻⁹ substitutions per site per generation, estimated for Galliformes (Ellegren 2007). These were selected because they represent empirical estimates from well-studied bird lineages and have been commonly applied in comparative genomic studies, and correspond to species with different life histories and genome characteristics, offering a conservative estimate of uncertainty in mutation rate. Both sets of results are reported in Table S3.

## Results

### Mitochondrial DNA divergence and population structure

The COI haplotype network revealed clear geographic structuring among populations (Figure 1B), with evidence of varying degrees of connectivity among regions. A mitochondrial lineage that was widely distributed throughout Lowland Bolivia, Northern and Central Argentina, and Uruguay, included about half of the *T. musculus* individuals (hereafter referred to as the Widespread mitochondrial lineage). In particular, it included a central and highly connected haplotype shared by individuals from multiple regions that may represent an ancestral or common haplotype. In contrast, populations from high altitude (> 3,400 m) in Bolivia (hereafter referred to as Highland Bolivia), Patagonia, and *T. cobbi* in the MFI formed distinct haplotype groups, each characterized by region-specific haplotypes more differentiated from the Widespread group, indicating that these lineages experience (or have experienced) some degree of isolation. Within Northern Argentina, a geographically localized haplotype lineage was found primarily in the eastern part of the region, where it co-occurs with the Widespread lineage and a few Patagonian individuals, suggesting a zone of lineage overlap. One sample from Highland Bolivia had mtDNA from the Widespread group, suggesting either recent movement of individuals among regions or a shared ancestral lineage. Most Patagonian individuals carried a unique haplotype, indicating very low mitochondrial variation in the region. This haplotype was also observed in Central and Northern Argentina, suggesting seasonal movements of Patagonian individuals. Only one individual differed from this pattern, carrying a second Patagonian haplotype primarily shared with individuals from Central Argentina, specifically from Mendoza.

The mean mitochondrial lineage divergence (K2P distance) ranged from 1.4% to 5.2% across population pairs (Figure 1D), corresponding to approximate divergence times of ∼0.7 to 2.5 million years based on an avian COI molecular clock of ∼2.1% per million years (Weir and Schluter 2008; Lavinia et al. 2016). The highest divergences were observed in comparisons involving *T. cobbi*, followed by the comparisons including Highland Bolivia, consistent with their degree of geographic isolation. In contrast, divergence among continental populations that are closer geographically (Patagonia, Widespread, and Northern Argentina) was comparatively lower, suggesting more recent or ongoing gene flow among these groups.

### Genome-wide population structure STRUCTURE analysis

The STRUCTURE analysis revealed clear geographic clustering among individuals. The Evanno method supported K = 3 as the most likely number of clusters, while the mean log probability of the data [LnP(Data)] indicated that K-values of four and five could capture additional finer-scale structure. The most geographically informative clustering solution was generated with a K value of four (Figure 1C), grouping *T. musculus* into three clusters: one including the Patagonian individuals (Patagonian cluster), one including the Bolivian high-altitude individuals (Highland Bolivia cluster) and a third group with the remaining continental birds (Widespread cluster) that included individuals from low altitude in Bolivia (Lowland Bolivia), Northern Argentina, Central Argentina and Uruguay, indicating higher levels of genetic connectivity in this broad area. The fourth cluster was restricted to *T. cobbi* individuals from the MFI, indicating strong differentiation in this insular-endemic species. All clusters were largely consistent with their mitochondrial counterparts, except for the absence of a nuclear cluster exclusive of Northern Argentina (the four individuals that belonged to this mitochondrial lineage were included in the Widespread nuclear genomic cluster). Notably, several individuals from the province of Mendoza in Central Argentina show extensive admixture between the Widespread and Patagonia clusters. Lowland Bolivian birds have admixed genomic content between the Highland Bolivian and Widespread clusters (to a lesser extent, some individuals from Northern Argentina also showed admixture between these two clusters, but with more content from the latter). This suggests extensive gene flow among continental *T. musculus* populations. Contrary to these birds with admixed genomic content, the finding of complete Patagonian genomic content in five individuals from Northern Argentina and four individuals from Central Argentina that also belonged to the Patagonian mitochondrial lineage (see Figure 1C) strongly suggests that these were Patagonian migrants (also supported by the fact that they were sampled in the non-reproductive season). Conversely to the high levels of admixture among *T. musculus* populations, there was virtually no presence of *T. cobbi* genomic content in the continental individuals (only seen in very low proportions), although some insular individuals did have continental genomic content. This suggest that there is overall low gene flow between *T. cobbi* and *T. musculus*, yet the presence of DNA from the latter in the MFI could also be due to the lack of lineage sorting after the islands were colonized.

It is worth mentioning that K = 3 shows most of the general patterns described for K = 4, but without a differentiated cluster in Highland Bolivia (i.e., the high Andean individuals were included in the Widespread cluster, Figure S1). To the contrary, increasing the number of clusters to K = 5 not only retrieved the Highland Bolivian cluster, but also generated a weak additional subgroup that corresponded to individuals from Mendonza in Central Argentina, suggesting that this area of contact between the Widespread and Patagonian clusters has some degree of genetic substructure (Figure S1).

### Principal Component Analysis

The PCA results were consistent with those from STRUCTURE, with Highland Bolivia and *T. cobbi* forming clearly distinct clusters along PC1 and PC2 (Figure 1E). Individuals from Lowland Bolivia, Northern and Central Argentina and Uruguay, and Patagonia were clustered closer together, although they were separated into the Widespread and the Patagonian clusters. Within the Widespread group, individuals from Lowland Bolivia, Northern and Central Argentina and Uruguay formed a diffuse cluster, with evident subtle geographic substructure: A) individuals from Northern Argentina clustered closer to Highland Bolivia, B) individuals from Central Argentina were more spread, with some of them placed within the Widespread cluster (but closer to the Patagonian individuals) and the ones from Mendoza in an intermediate position between the Widespread and the Patagonia clusters, and C) two lowland Bolivian individuals were clearly placed within the Widespread cluster but the other two had an intermediate position between the Widespread and the Highland Bolivia clusters. Finally, the Patagonian cluster consisted of individuals from Patagonia, along with some from Northern and Central Argentina, which may be migrants captured in their wintering grounds as mentioned above.

### Fine-scale population structure

The fineRADstructure analysis further confirmed genetic differentiation among major geographic regions, revealing an initial split between northern and southern populations, which was subsequently resolved into four distinct clusters (Figure 1F). The northern group comprised individuals from Highland Bolivia and the Widespread cluster, which included samples from Lowland Bolivia, Northern and Central Argentina, and Uruguay. Within this group, a latitudinal genetic gradient was again apparent. The southern group was further divided into two clusters: *T. cobbi* from the MFI and Patagonia. Again, the individuals from Mendoza (the group showing some differentiation in the STRUCTURE analysis at K = 5) appear to be related to individuals from the Widespread and the Patagonian groups. The MFI and Highland Bolivia exhibited the highest levels of differentiation.

### Phylogenetic inference

Phylogenetic analyses of mitochondrial DNA using IQ-TREE and RAxML-NG show strong concordance, consistently recovering a clear north–south geographic split among samples (Figure 2A, S2A). Both trees reveal a deep division separating a northern clade—comprising individuals from Central Argentina, Northern Argentina, Lowland and Highland Bolivia, and Uruguay—from a southern clade that includes samples from *T. cobbi* and Patagonia along with individuals from Central and Northern Argentina that may be Patagonian migrants. We found additional structure within each of these larger clades as in the previous analyses. The first division within the northern clade is weakly supported, but subsequent nodes show higher support, including a Widespread clade and two more defined clades: one composed of individuals from Northern Argentina, and another consisting primarily of Highland Bolivia individuals. Notably, one individual from Highland Bolivia is found within the Widespread group. Within the southern clade, one well-supported group consists of individuals from the MFI, forming a distinct monophyletic clade. The second group includes two subclades with intermediate support: one containing individuals from Central Argentina, specifically from Mendoza, including one Patagonian sample, and the other comprising individuals from Patagonia as well as the presumably migrants from Northern and Central Argentina.

**Figure 2.**
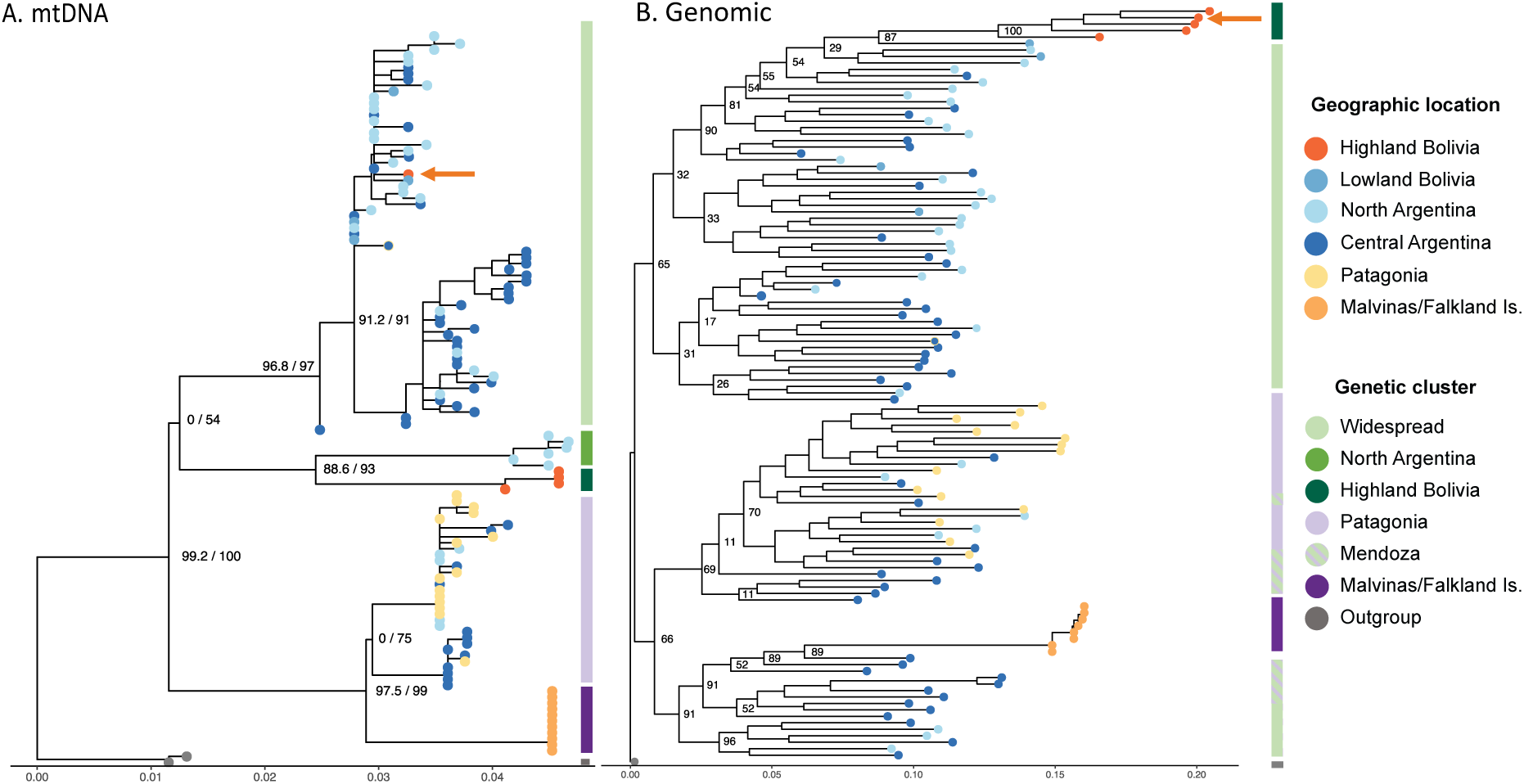
Phylogenetic relationships among *Troglodytes muculus* and *T. cobbi* individuals. **(A)** Phylogenetic tree based on mitochondrial COI (cytochrome oxidase I) sequences from 121 individuals. **(B)** Phylogenetic tree based on genome-wide SNP data from 116 individuals. Both trees were inferred using IQ-TREE. Tip colors indicate geographic origin, and the vertical color bars to the right of each tree denote genetic clusters (mtDNA and nuclear genomic DNA, respectively). The x-axis represents the number of substitutions per site, reflecting genetic divergence among haplotypes. The orange arrow highlights the case of mito-nuclear discordance also illustrated in Figure 1.

Support across phylogenetic trees based on genome-wide SNPs is generally low, with the notable exceptions of strongly supported Highland Bolivia and MFI clades, consistent with other analyses (Figure 2B, S2B). The low-resolution limits confidence in finer-scale relationships and suggests these phylogenetic relationships could be blurred through gene flow. Comparison of mitochondrial and nuclear phylogenies shows broad agreement in major population divisions, including the north–south split and the distinct clades corresponding to Highland Bolivia and *T. cobbi* from the MFI. At finer scales, topologies differ, with mitochondrial trees generally exhibiting clearer structure than nuclear SNP trees, which show lower resolution and weaker support.

### Few instances of mito-nuclear discordance

Overall, we found broad concordance between mitochondrial and nuclear genetic structure, with both types of genetic markers supporting the same population divisions across the species’ range (Figure 1). This agreement suggests that historical isolation shaped both genomes similarly. However, we observed a few instances of mito-nuclear discordance between geographically proximate populations, consistent with recent or ongoing gene flow or incomplete lineage sorting. A mitochondrial lineage restricted to northeastern Argentina lacked corresponding nuclear differentiation, with all individuals assigned to the Widespread genomic group. We also detected a Patagonian mitochondrial haplotype associated with the Widespread nuclear genome. Lastly, one individual from Highland Bolivia possessed a Widespread mitochondrial haplotype, while its nuclear genome placed it within the Highland Bolivia clade (orange arrow example in Figure 1A, 1B, 1E, 2, S2). These exceptions likely reflect gene flow, incomplete lineage sorting, or potential sex-biased dispersal, which can differentially affect mitochondrial and nuclear patterns, highlighting the importance of integrating multiple genomic markers when reconstructing population history.

### Isolation by distance and effective migration and diversity surfaces (EEMS)

We detected a significant pattern of isolation by distance (IBD) across the sampled portion of the species’ range, with a moderate positive correlation between genetic and geographic distances (Mantel r = 0.383, *p* = 0.001; Figure 3A), which was also supported by MRM (R² = 0.147, p = 0.001). These results are consistent with a model of gene flow among neighboring populations. To explore genetic differentiation beyond what is expected under IBD, we conducted an effective migration surface (EEMS) analysis. EEMS revealed pronounced spatial heterogeneity in genetic diversity across the species’ range, highlighting a hotspot in central Argentina, which gradually decreases toward the south and northwest. Lower diversity rates were observed in peripheral regions such as southern Patagonia and for *T. cobbi*, suggesting potential demographic bottlenecks or founder effects leading to reduced effective population sizes in these areas (Figure 3B). Consistently, the mean migration rates show elevated gene flow in a central corridor encompassing parts of Northern and Central Argentina, while migration rates were markedly reduced across the Bolivian highlands, southern Patagonia and for *T. cobbi* in the MFI. These findings suggest the presence of migratory corridors, and highlight the 400 km of sea separating continental Patagonia from the MFI and the Andes as strong barriers to gene flow (Figure 3C).

**Figure 3.**
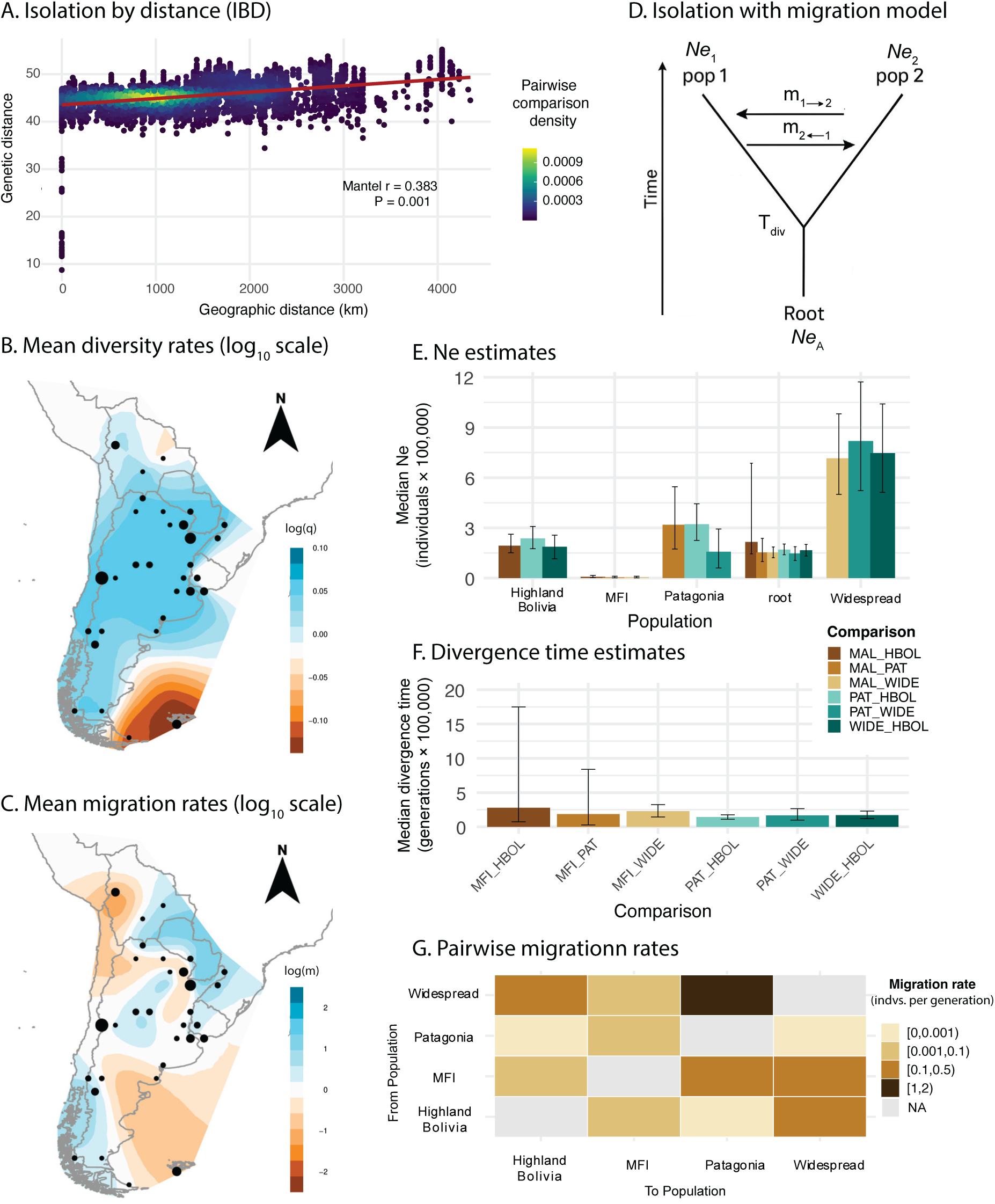
Spatial patterns of genetic diversity, gene flow and demographic reconstruction. **A)** Isolation by distance (IBD) in the Southern and Cobb’s Wrens. Scatterplot of pairwise genetic distance versus geographic distance (in kilometers) for all individuals, based on genome-wide unlinked SNP data. Each point represents a pairwise comparison and is colored by local point density, highlighting regions of the plot with higher sampling intensity. The color scale reflects relative point density estimated via two-dimensional kernel density smoothing, with warmer colors indicating more densely clustered comparisons. The solid line shows the linear regression fit, consistent with a pattern of isolation by distance across the species’ range. **B)** Mean diversity rates (log₁₀ scale of α) across the species range inferred using EEMS, showing areas of elevated genetic diversity in central Argentina and reduced diversity in peripheral regions such as southern Patagonia and the Malvinas/Falkland Islands (MFI). **C)** Mean migration rates (log₁₀ scale) inferred by EEMS, with high gene flow corridors in central Argentina and reduced migration across the Andes and the MFI. **D)** Schematic representation of the isolation-with-migration (IM) model used in the G-PhoCS analyses. Each pairwise model includes two extant populations (pop1 and pop2) and one ancestral population (root). Parameters estimated include the effective population sizes (Ne_1_, Ne_2_, and ancestral Ne_A_), the divergence time (Tdiv), and bidirectional migration rates (m1→2, m2→1) after divergence. All population comparisons were run in a pairwise approach following this framework. **E)** Median effective population size (*Ne*) estimates (with 95% HPD intervals) from G-PhoCS for each regional population and the root, showing variation in effective population sizes across populations. **F)** Median divergence time estimates (with 95% HPD intervals) between population pairs inferred by G-PhoCS, with deeper divergence involving the MFI population. Population abbreviations: HBOL—Highland Bolivia; WIDE—Widespread; MFI—Malvinas/Falkland Islands; PAT—Patagonia. The results shown for *Ne* and divergence time estimates were scaled to individuals and generations using the mutation rate of 4.6 x 10^-9^ (Smeds et al. 2016). **G)** Pairwise migration rate estimates among populations from G-PhoCS expressed as numbers of migrants per generation, with rates binned into categories and visualized as a heatmap.

### Admixed origin of individuals from the province of Mendoza

Our results show that individuals from Mendoza are intermediate between the Patagonian and Widespread nuclear clusters (Figure 1) yet also show weak signals of having their own genetic identity (Figure S1). These patterns are replicated when we analyze these focal populations in the absence of the remaining samples in the dataset (Figure S3). We decided to explore the genetic composition of these intermediate individuals using triangle plots, with the caveat that we found few differentiated markers between the parental lineages (Patagonia and Widespread) to begin with, leading to generally low resolution and most parental individuals from either population appearing as intermediate themselves (Figure S3D). When using a threshold of allele frequency difference of 0.4, individuals from Mendoza were all distant from the triangle extremes, frequently located around a hybrid index of 0.5, and exhibited relatively high heterozygosity, consistent with early-generation hybrids generated after the secondary contact between lineages.

### Demographic inference

We used an isolation-with-migration (IM) model to infer divergence times, effective population sizes (*Ne*), and gene flow among populations (Figure 3D). Because the assumption of mutation rate greatly impacts the absolute values of parameter estimates, we focus on relative differences among groups. To reflect the uncertainty in demographic scaling we also scaled our estimates using two mutation rates reported in the literature for birds (Table S3). Estimates of *Ne* showed substantial variation across groups. The Widespread population has the largest Ne values, whereas Highland Bolivia and *T. cobbi* had notably lower estimates (consistent with estimates of genetic diversity, Figure 3B). *T. cobbi* had the lowest Ne, with estimates an order of magnitude lower than continental groups, suggesting a strong historical bottleneck likely generated by a founder effect followed by prolonged isolation (Figure 3E, Table S3). Divergence time estimates indicated recent splits among most population pairs and generally followed what would be expected based on the phylogenetic relationship among populations (Figure 2). The lowest divergence times were found between geographically contiguous populations such as Widespread and Highland Bolivia, as well as between Patagonia and the Widespread group. In contrast, divergence involving *T. cobbi* with Highland Bolivia and the Widespread groups, showed higher median values. Notably, the divergence time in models including *T. cobbi* converged poorly in all comparisons and show high uncertainty, suggesting challenges in estimating divergence parameters for this group (Figure 3F, Table S3). Pairwise migration rate estimates revealed asymmetric and heterogeneous gene flow between populations. However, estimates of migration direction from G-PhoCS should be interpreted cautiously, as the method has reduced power to recover directionality (Gronau et al. 2011). Migration involving *T. cobbi* was generally lower, consistent with its lower diversity, higher divergence estimates, and greater geographic isolation. The low but detectable levels of migration between the MFI and the continent might reflect shared genomic variants that have not yet sorted after the colonization of the islands rather than actual gene flow. These patterns reflect both historical isolation in the MFI and contemporary continental gene flow dynamics among regional populations (Figure 3G, Table S3).

## Discussion

Our analysis of spatial population structure combining mitochondrial DNA and genome-wide SNP data revealed geographically distinct lineages with a complex evolutionary history within the Southern House Wren’s and Cobb’s Wren’s ranges in southern South America, despite a relatively continuous continental distribution of the Southern House Wren. Consistently across analyses and marker types, we observe: 1) a latitudinal split between northern populations of *T. musculus* (including Bolivia, Northern and Central Argentina, and Uruguay) and the southern populations of *T. musculus* and *T. cobbi* (Patagonia and the MFI, respectively), 2) higher genetic differentiation in the insular species (*T. cobbi*), with negligible gene flow with the continent, and 3) a contrasting pattern within the continent, with notorious gene flow in the secondary contact zones between the northern and southern lineages, although with reduced gene flow between the Andean and lowland populations. We discuss these findings below, emphasizing the putative role of Pleistocene glaciations in directly (through vicariance) or indirectly (through the effects of climate on habitat dynamics) shaping the evolutionary history of this species complex in southern South America.

Notably, both mitochondrial and nuclear data consistently identify the deepest divergence within southern South America as occurring between the southern lineage (comprising the Patagonian populations of *T. musculus* and *T. cobbi*) and the northern *T. musculus* populations. We use the timing of the split between the Widespread and Patagonian clades as a proxy for this north–south divergence (as we only estimated divergence in G-PhoCS using current and not ancestral populations). Additionally, both comparisons show nearly identical mitochondrial divergence levels (mean genetic distances of 0.036–0.037, respectively), suggesting they capture the same major phylogeographic break. The timing of this event remains uncertain and depends on the assumed mutation rate and the type of genetic marker used. The mtDNA estimate places the divergence at between 1.4 and 1.7 million years ago (based on D_a_ and K2P distances, respectively), whereas nuclear DNA suggests a more recent split. The demographic estimates based on nuclear data span approximately 160 thousand to 600 thousand years ago, depending on the mutation rate applied (1.2 × 10⁻⁹ substitutions per site per million years obtained for Galliformes vs. 4.6 × 10⁻⁹ substitutions per site per generation, estimated for flycatchers). Moreover, there is also considerable uncertainty in many divergence time estimates, with broad 95% credible intervals (Table S3). Nuclear mutation rates are generally more uncertain than mtDNA rates, which are often better calibrated in birds, therefore, mtDNA may provide a more reliable point estimate (Allio et al. 2017). In addition, because nuclear differences between populations or lineages can be affected by gene flow, which we have shown that in fact occurs among the continental populations of *T. musculus*, divergence times might be underestimated by nuclear DNA (Lavinia et al. 2019). Despite these uncertainties, the estimated divergence broadly coincides with the Great Patagonian Glaciation, a period encompassing at least three major glacial advances between approximately 2.6 and 0.78 million years ago (Griffing et al. 2022). The most extensive of these, occurred around 1.17 ± 0.14 Ma, and represents the maximum glacial extent in the region, during which ice expanded across non-Andean Patagonia, reaching as far as the Atlantic coast in the southernmost portion of the continent (Rabassa and Coronato 2009; Rabassa et al. 2011; Griffing et al. 2022). The presence of these glaciers isolated plants and vertebrates, which diversified in multiple ice-free refugia throughout Patagonia (Nuñez et al. 2011; Sersic et al. 2011; Marín et al. 2013; Sánchez et al. 2024), including also various recently described cases of intraspecific diversification in birds (Acosta et al. 2021; Bukowski et al. 2024; Balza et al. 2025; Lavinia et al. 2025), suggesting that this could also be the case for the Patagonian lineage of *T. musculus*.

Shortly after the initial divergence between the two main *T. musculus* lineages, further intra-lineage diversification occurred. In the southern lineage, Patagonian individuals likely colonized the MFI, where this newly formed insular population became isolated from the continent and diverged with virtually no gene flow. This colonization may have been facilitated by a shorter distance between the islands and the continent during glacial periods compared to the current ∼400 km oceanic separation, or even the presence of a land bridge connecting them (Ponce et al. 2011). In fact, the biota of these islands, including their avifauna, has been considered to be derived from that of southern Patagonia and Tierra del Fuego (McDowall 2005; Morrone and Posadas 2005). This scenario is supported by the sister relationship recovered between the Patagonian lineage of *T. musculus* and *T. cobbi* by previous mitochondrial results (Campagna et al. 2012) and by both the mitochondrial and nuclear data in this study. In addition, this insular population is the least diverse both in its mitochondrial DNA (a unique haplotype was shared among all sampled individuals) and in its nuclear genomic DNA, a result consistent with a founder effect. Finally, both DNA sources coincide in placing this event around the mid-Pleistocene, with mtDNA suggesting the divergence approximately 0.89 - 1.4 million years ago (based on D_a_ and K2P distances, respectively), and the nuclear estimates placing the split between ∼200-700 thousand years ago, depending on the mutation rate used. Again, these estimates are associated with broad 95% credible intervals, with the upper limit of the nuclear estimate using the flycatcher rate approaching the lower bound of the mtDNA divergence, and both mtDNA estimates falling within the interval when applying the Galliformes rate. Irrespective of the differences between markers, these timings coincide with the scenario of a colonization of the MFI from Patagonia and its posterior isolation shortly after the initial divergence between the two main lineages.

Due to the absence of gene flow between the MFI and Patagonia, the insular population experienced a process of divergence that resulted in speciation. *T. cobbi* has been in fact recognized as a distinct species for several years as it has been shown to possess not only mitochondrial differences, but also certain diverged nuclear loci and morphological and vocal differentiation (Campagna et al. 2012; Chesser et al. 2024). Our results confirm this interpretation, as *T. cobbi* was shown to have a clearly differentiated genome from the continental populations, being in fact the most differentiated in the entire southern portion of South America. This pronounced distinctiveness is consistent with the divergence of the MFI populations in other bird species, in which the insular isolation has led to the emergence of endemic or independently evolving lineages with various levels of differentiation—including *Cistothorus platensis*, *Melanodera melanodera*, *Turdus falcklandii*, *Tachyeres brachypterus*, *Chloephaga rubidiceps*, and *Chloephaga picta* (Campagna et al. 2012, 2019; Kopuchian et al. 2016). On the other hand, this result is also congruent with recent findings in the House wren species complex, in which various insular forms have been shown to be divergent and were split as different species (*T. martinicensis*, *T. mesoleucus. T. beani, T. musicus and T. grenadensis* from Dominica, St. Lucia, Cozumel, St. Vincent, and Grenada islands, respectively; (Klicka et al. 2023; Chesser et al. 2024).

This sister relationship between *T. cobbi* and the Patagonian lineage of *T. musculus* implies that the latter species is paraphyletic. However, paraphyly does not necessarily undermine its species status, as many recognized species are in fact paraphyletic (Funk and Omland 2003; McKay and Zink 2010), particularly several cases involving insular diversification (Emerson 2002). Paraphyly is especially common when divergence is recent and incomplete lineage sorting persists (McKay and Zink 2010), and is expected when the new species originates from a species with a wide distribution such as the Southern House Wren. Consistently, the same situation occurs with the other aforementioned insular species of the House Wren complex.

A split concurrent in timing with that of the MFI/Patagonian population occurred in the northern lineage, where the Andean populations of Bolivia became isolated from the Widespread, low altitude lineage at around 1.5-1.7 million years ago according to mtDNA divergence (D_a_ and K2P, respectively) and between 170-650 thousand years ago using nuclear DNA. In this case, the nuclear estimates are considerably more recent and their 95 credible intervals are narrower, so the mtDNA and nuclear estimates are farther apart. This lineage, which distinctiveness is congruent with the diversification produced by glacial cycles in high-altitude Andean birds (Weir 2006; Jetz et al. 2012; Hazzi et al. 2018), might extend to southern unsampled Andean localities in Argentina and to the north, and coincide with the Andean populations of *T. musculus* detected by Klicka et al. (2023) in central and southern Peru. This Andean populations are the most differentiated within the southern distribution of *T. musculus* but, unlike the insular *T. cobbi*, they show some degree of gene flow with other continental populations and the presence of admixed genomic content in the birds of the zone of secondary contact with the Widespread lineage, particularly in lowland Bolivia and to a lesser extent in northern Argentina.

Considerable gene flow has been detected among continental populations, particularly in the area of secondary contact between the Widespread and Patagonian lineages across central Argentina, where admixed genomic content is prevalent. This is especially notable in west-central Argentina, in the Andean slopes of Mendoza province, where around half of the individuals have the Widespread mitochondrial lineage, the other half have the Patagonian mitochondrial lineage and all of them have around 50% content of each of these nuclear genomic clusters. This pattern, where both mitochondrial lineages are present and the individuals possess varying degrees of nuclear admixture in central Argentina, has also been found in other widespread birds with secondary contact after the isolation of Patagonian populations in glacial periods, including *Zonotrichia capensis* (Campagna et al. 2014) and *Vanellus chilensis* (Bukowski et al. 2024).

Interestingly, a closer look at the west-central population in Mendoza, including analyses performed solely with the Widespread and Patagonian genomic clusters, revealed some substructure within this locality. In the PCA including all samples, Mendoza individuals fall roughly in the middle between the Widespread and Patagonian clusters, consistent with an initial admixture between these lineages. However, variation along PC2 indicates additional differentiation within Mendoza, suggesting that after the initial admixture event, this population may have experienced some degree of isolation and divergence, and is now somewhat distinct from the parental clusters. This pattern is further supported by other analyses, such as the emergence of a distinct genomic cluster for the Mendoza population at K=5 and K=3 in the STRUCTURE analyses using the complete dataset and the three-population subset, respectively.

A subset of individuals from central and northern Argentina cluster entirely with Patagonian individuals based on nuclear DNA, instead of showing intermediate genomic content between the Widespread and Patagonian clusters, and are placed within the mitochondrial Patagonian lineage. All these individuals were sampled in the non-breeding season (between April and October) and therefore appear to be Patagonian migrants that moved northwards during Autumn and Winter. Although migration in southern South America has been understudied compared to the Northern Hemisphere, several Patagonian populations of other species are believed to migrate in the non-breeding season, and this has been recently confirmed for some of them (e.g., Jara et al. 2024; Lisovski et al. 2025). Consistent with this interpretation, in the Northern House Wren (*T. aedon*), the northern *T. a. aedon* and *T. a. parkmanii* undertake seasonal migrations (Johnson 2024), though the extent of their wintering ranges remains unclear (Anderson et al. 2025). Further ecological and tracking studies are needed to confirm this hypothesis.

Mitochondrial and nuclear patterns were generally concordant, yet we observed instances of mito-nuclear discordance, as was also the case in the previous study of this species complex performed by Klicka et al. (2023). We found possible cases of mitochondrial introgression, such as the individual sampled in the Bolivian Highlands with the typical genomic content of this cluster but a Widespread mitochondrial haplotype. However, the most notorious discordance between the mitochondrial and nuclear results was the presence of a mitochondrial lineage represented by a few individuals from northern Argentina that did not have any genomic correlate (i.e., these individuals fell within the widespread genomic cluster and could not be distinguished from the rest of the representatives of this cluster). A broader geographic sampling northwards would be needed to determine whether this is a local mitochondrial lineage or these individuals were the southernmost representatives of a lineage that extends to the north into Brazil or Paraguay.

This type of discordance between mitochondrial and nuclear markers is well documented and has been attributed to several processes (Funk and Omland 2003; Zink and Barrowclough 2008; Toews and Brelsford 2012). For instance, because the mitochondrial genome is haploid and maternally inherited, it has a fourfold smaller effective population size than nuclear DNA, allowing it to sort lineages more rapidly (Hudson and Turelli 2003; Zink and Barrowclough 2008). This can lead to discrepancies if mtDNA has already reached fixation while the nuclear genome retains ancestral variation. Additionally, sex-biased dispersal (e.g., female-mediated gene flow) or asymmetric introgression following hybridization can introduce mtDNA haplotypes across population boundaries without similar patterns in the nuclear genome (Funk and Omland 2003; Rheindt and Edwards 2011). Moreover, high levels of gene flow could have homogenized the nuclear DNA while mtDNA differences will persist until one of the mitochondrial haplotypes is lost by drift. Finally, natural selection on mitochondrial genes may also drive mismatched patterns if selection varies geographically (Bazin et al. 2006; Meiklejohn et al. 2007; Irwin 2012; Morales et al. 2018).

## Final remarks

In sum, our study shows that glacial cycles were likely the main drivers of the diversification of the Southern House Wren complex in southern South America causing a series of splits coincident with the Great Patagonian Glaciation. These included the differentiation of Patagonian populations, and shortly afterwards the colonization and isolation in the MFI and the divergence of an Andean lineage. After glaciers retreated, continental populations entered into secondary contact, resuming gene flow and generating a broad area of admixed genomic content in central Argentina, and to a lesser extent in northern Argentina and lowland Bolivia. To the contrary, the sea-level rise caused by the end of this glacial period increased the isolation of the MFI, ceasing the exchange of genetic material between the continental populations and their insular counterpart and promoting the differentiation, and eventually speciation, of the population of the archipelago.

## Author contributions

Conceptualization: DAL, CK, PLT, LC

Methodology: MR, DAL, LC

Investigation: MR, DAL, CK, PAF, PLT, LC.

Visualization: MR

Writing—original draft: MR, LC, DAL

Writing—review & editing: MR, DAL, CK, PLT, LC

## Acknowledgements

This project was funded by NSF DEB-2232929 (to LC) CONICET (to CK, PAF, PLT and DAL), Agencia Nacional de Promoción de la Investigación, el Desarrollo Tecnológico y la Innovación (to PAF, PLT and DAL), Richard Lounsbery Foundation (to PLT and DAL) and the International Barcode of Life project.

## Data availability Statement

All mitochondrial and genomic sequence data supporting this study are available in Figshare at: https://doi.org/10.6084/m9.figshare.29226047. Genomic data have been archived in GenBank (BioProject ID PRJNAXXXXX). All the Accession numbers are specified in the Table S2.

