## Supplementary Material for "Continental diversification and insular speciation in a widespread passerine (*Troglodytes musculus*) in southern South America"

**Figure S3. Genomic variation and evidence of admixture in the Mendoza population**... **17**

***Supplementary Tables***

**Table S1. General information for Troglodytes musculus/cobbi samples across southern South America.** The table includes the population of origin, collection date, the geographic coordinates (Latitude and Longitude) of sampling locations and the Accession numbers (Genbank for COI and NCBI for ddRAD data). Museum/collections abbreviations: Museo Argentino de Ciencias Naturales “Bernardino Rivadavia” (MACN); Burke Museum of Natural History and Culture, University of Washington (UWBM); Queen’s University Molecular Ecology Laboratory collection (QUMEL).

| Sample ID | Population | Collection date (dd/mm/yyyy) | Lat | Long | Accession number COI | Accession number ddRAD |
|---|---|---|---|---|---|---|
| MACN-Or-ct 1016 | Northern Argentina | 3/11/05 | -23.91 | -65.48 | FJ028458 | TBD |
| MACN-Or-ct 1460 | Northern Argentina | 28/04/2006 | -27.55 | -58.68 | JQ943367 | TBD |
| MACN-Or-ct 1475 | Northern Argentina | 29/04/2006 | -27.55 | -58.68 | FJ028462 | TBD |
| MACN-Or-ct 1631 | Northern Argentina | 10/5/06 | -27.55 | -58.68 | JQ943368 | TBD |
| MACN-Or-ct 1710 | Northern Argentina | 31/05/2006 | -27.58 | -58.65 | JQ943369 | TBD |
| MACN-Or-ct 1860 | Northern Argentina | 25/10/2006 | -27.55 | -58.68 | FJ028471 | TBD |
| MACN-Or-ct 1895 | Central Argentina | Feb-07 | -34.55 | -58.5 | FJ028473 | TBD |
| MACN-Or-ct 1902 | Central Argentina | 22/11/2006 | -32.78 | -59.91 | FJ028469 | TBD |
| MACN-Or-ct 1903 | Central Argentina | 22/11/2006 | -32.78 | -59.91 | FJ028474 | TBD |
| MACN-Or-ct 2333 | Patagonia | 30/12/2006 | -54.51 | -67.19 | FJ028470 | TBD |
| MACN-Or-ct 2412 | Northern Argentina | Jun-07 | -27.57 | -58.69 | JQ943370 | TBD |
| MACN-Or-ct 2586 | Patagonia | 30/10/2006 | -41.79 | -71.56 | TBD | TBD |
| MACN-Or-ct 2599 | Patagonia | 31/10/2006 | -41.79 | -71.56 | FJ028475 | TBD |
| MACN-Or-ct 2624 | Patagonia | 2/11/06 | -41.79 | -71.56 | JQ943371 | TBD |
| MACN-Or-ct 2666 | Patagonia | 5/11/06 | -41.79 | -71.56 | FJ028466 | TBD |
| MACN-Or-ct 2679 | Patagonia | 8/11/06 | -41.95 | -71.4 | FJ028464 | TBD |
| MACN-Or-ct 2680 | Patagonia | 8/11/06 | -41.95 | -71.4 | FJ028465 | TBD |

| | | | | | | |
|---|---|---|---|---|---|---|
| MACN-Or-ct 2846 | Northern Argentina | 7/12/06 | -25.68 | -54.44 | FJ028468 | TBD |
| MACN-Or-ct 286 | Patagonia | Feb-05 | -51.72 | -70.17 | FJ028463 | TBD |
| MACN-Or-ct 2893 | Northern Argentina | 11/12/06 | -25.68 | -54.44 | FJ028467 | TBD |
| MACN-Or-ct 3168 | Northern Argentina | 20/07/2007 | -25.12 | -58.17 | FJ028476 | TBD |
| MACN-Or-ct 3196 | Northern Argentina | 23/07/2007 | -25.12 | -58.17 | FJ028472 | TBD |
| MACN-Or-ct 3335 | Central Argentina | 12/11/05 | -34.46 | -58.55 | JQ943372 | TBD |
| MACN-Or-ct 3546 | Northern Argentina | 22/11/2007 | -25.68 | -54.44 | JQ943373 | TBD |
| MACN-Or-ct 3636 | Central Argentina | 31/12/2007 | -34.62 | -58.41 | JQ943374 | TBD |
| MACN-Or-ct 3759 | Patagonia | 21/02/2007 | -41.13 | -71.31 | JQ943375 | TBD |
| MACN-Or-ct 3884 | Lowland Bolivia | 27/05/2008 | -15.66 | -67.46 | JQ943364 | TBD |
| MACN-Or-ct 4100 | Central Argentina | 19/01/2009 | -34.64 | -58.46 | TBD | TBD |
| MACN-Or-ct 4162 | Northern Argentina | 28/08/2008 | -27.32 | -58.95 | HQ569914 | TBD |
| MACN-Or-ct 4179 | Northern Argentina | 29/08/2008 | -27.32 | -58.95 | HQ569923 | TBD |
| MACN-Or-ct 4204 | Northern Argentina | 30/08/2008 | -27.32 | -58.95 | HM396287 | TBD |
| MACN-Or-ct 4248 | Central Argentina | 22/10/2008 | -35.13 | -57.39 | HM396072 | TBD |
| MACN-Or-ct 4271 | Central Argentina | 23/10/2008 | -35.13 | -57.39 | HM396082 | TBD |
| MACN-Or-ct 4344 | Central Argentina | 15/11/2008 | -31.89 | -58.24 | JQ943366 | TBD |
| MACN-Or-ct 4345 | Central Argentina | 15/11/2008 | -31.89 | -58.24 | JQ943376 | TBD |
| MACN-Or-ct 4355 | Central Argentina | 16/11/2008 | -31.89 | -58.24 | JQ943360 | TBD |
| MACN-Or-ct 4516 | Northern Argentina | 1/2/09 | -26.81 | -59.61 | HQ569978 | TBD |
| MACN-Or-ct 4560 | Patagonia | 15/01/2008 | -40.16 | -71.39 | JQ943361 | TBD |
| MACN-Or-ct 4561 | Patagonia | 12/12/08 | -40.16 | -71.4 | JQ943362 | TBD |
| MACN-Or-ct 4563 | Central Argentina | 5/5/09 | -35.13 | -57.39 | HM396111 | TBD |
| MACN-Or-ct 4567 | Central Argentina | 5/5/09 | -35.14 | -57.39 | HM396114 | TBD |
| MACN-Or-ct 4579 | Central Argentina | 6/5/09 | -35.13 | -57.39 | HM905953 | TBD |
| MACN-Or-ct 4663 | Lowland Bolivia | 29/06/2009 | -15.46 | -67.48 | JQ943365 | TBD |
| MACN-Or-ct 4785 | Northern Argentina | 5/9/09 | -26.79 | -59.62 | HM396137 | TBD |
| MACN-Or-ct 4793 | Northern Argentina | 6/9/09 | -26.79 | -59.62 | JQ943363 | TBD |
| MACN-Or-ct 4809 | Northern Argentina | 6/9/09 | -26.79 | -59.62 | HM396146 | TBD |

| | | | | | | |
|---|---|---|---|---|---|---|
| MACN-Or-ct 4811 | Northern Argentina | 7/9/09 | -26.79 | -59.62 | HM396148 | TBD |
| MACN-Or-ct 5016 | Northern Argentina | 17/10/2009 | -28.02 | -58.02 | HM396216 | TBD |
| MACN-Or-ct 5063 | Northern Argentina | 19/10/2009 | -28.02 | -58.02 | HM396247 | TBD |
| MACN-Or-ct 5064 | Northern Argentina | 20/10/2009 | -28.02 | -58.02 | TBD | TBD |
| MACN-Or-ct 5077 | Northern Argentina | 20/10/2009 | -28.02 | -58.02 | HM396253 | TBD |
| MACN-Or-ct 5101 | Central Argentina | 15/12/2009 | -38.53 | -61.89 | HM396262 | TBD |
| MACN-Or-ct 5125 | Central Argentina | 18/12/2009 | -38.03 | -62.19 | HM396265 | TBD |
| MACN-Or-ct 5326 | Central Argentina | Sep-09 | -30.89 | -63 | TBD | TBD |
| MACN-Or-ct 5327 | Central Argentina | Sep-09 | -30.89 | -63 | TBD | TBD |
| MACN-Or-ct 5336 | Central Argentina | 23/02/2010 | -34.05 | -67.95 | TBD | TBD |
| MACN-Or-ct 5384 | Northern Argentina | 28/06/2010 | -24.31 | -61.81 | TBD | TBD |
| MACN-Or-ct 55 | Patagonia | 12/11/03 | -40.58 | -62.88 | FJ028461 | TBD |
| MACN-Or-ct 5669 | Northern Argentina | 6/8/10 | -23.75 | -64.85 | TBD | TBD |
| MACN-Or-ct 5684 | Northern Argentina | 7/8/10 | -23.75 | -64.85 | TBD | TBD |
| MACN-Or-ct 5896 | Lowland Bolivia | 22/10/2010 | -17.44 | -62.17 | TBD | TBD |
| MACN-Or-ct 599 | Patagonia | 25/01/2005 | -41.16 | -70.41 | FJ028460 | TBD |
| MACN-Or-ct 6112 | Central Argentina | 10/6/10 | -30.89 | -63 | TBD | TBD |
| MACN-Or-ct 6126 | Central Argentina | 16/04/2010 | -30.89 | -63 | TBD | TBD |
| MACN-Or-ct 6178 | Northern Argentina | 29/05/2011 | -25.1 | -58.15 | TBD | TBD |
| MACN-Or-ct 6279 | Lowland Bolivia | 7/10/11 | -19.39 | -64.09 | TBD | TBD |
| MACN-Or-ct 6386 | Patagonia | 16/02/2011 | -50.33 | -73.24 | TBD | TBD |
| MACN-Or-ct 6701 | Highland Bolivia | 14/07/2011 | -16.54 | -68.06 | TBD | TBD |
| MACN-Or-ct 6702 | Highland Bolivia | 14/07/2011 | -16.54 | -68.06 | TBD | TBD |
| MACN-Or-ct 6720 | Highland Bolivia | 16/08/2011 | -16.54 | -68.06 | TBD | TBD |
| MACN-Or-ct 6721 | Highland Bolivia | 16/08/2011 | -16.54 | -68.06 | TBD | TBD |
| MACN-Or-ct 6722 | Highland Bolivia | 16/08/2011 | -16.54 | -68.06 | TBD | TBD |
| MACN-Or-ct 6928 | Northern Argentina | 10/10/12 | -27.44 | -54.94 | TBD | TBD |
| MACN-Or-ct 6937 | Northern Argentina | 10/10/12 | -27.44 | -54.94 | TBD | TBD |
| MACN-Or-ct 695 | Patagonia | 25/01/2005 | -41.16 | -70.41 | FJ028459 | TBD |

| | | | | | | |
|---|---|---|---|---|---|---|
| MACN-Or-ct 7022 | Central Argentina | 21/11/2012 | -31.69 | -64.85 | TBD | TBD |
| MACN-Or-ct 7065 | Central Argentina | 27/11/2012 | -31.73 | -64.85 | TBD | TBD |
| MACN-Or-ct 7528 | Northern Argentina | 11/2/13 | -27.44 | -54.94 | TBD | TBD |
| MACN-Or-ct 7554 | Central Argentina | 9/4/13 | -34.6 | -58.44 | TBD | TBD |
| MACN-Or-ct 7577 | Central Argentina | 22/11/2013 | -35.34 | -57.37 | TBD | TBD |
| MACN-Or-ct 7629 | Central Argentina | Dec-13 | -34.6 | -58.43 | TBD | TBD |
| MACN-Or-ct 7672 | Patagonia | 22/01/2012 | -50.33 | -73 | TBD | TBD |
| MACN-Or-ct 7702 | Central Argentina | 5/12/14 | -34.6 | -58.44 | TBD | TBD |
| MACN-Or-ct 7736 | Northern Argentina | 26/01/2015 | -28.02 | -58.04 | TBD | TBD |
| MACN-Or-ct 7849 | Central Argentina | 21/03/2015 | -31.69 | -64.85 | TBD | TBD |
| MACN-Or-ct 7872 | Central Argentina | 23/03/2015 | -31.69 | -64.85 | TBD | TBD |
| MACN-Or-ct 8269 | Northern Argentina | 24/10/2015 | -25.92 | -61.71 | TBD | TBD |
| MACN-Or-ct 8277 | Central Argentina | 6/9/15 | -34.82 | -57.96 | TBD | TBD |
| MACN-Or-ct 8674 | Uruguay | 29/10/1997 | -34.88 | -56.12 | TBD | TBD |
| MACN-Or-ct 8772 | Uruguay | 23/06/2001 | -34.78 | -55.59 | TBD | TBD |
| MACN-Or-ct 8916 | Uruguay | 9/1/06 | -32.48 | -53.75 | TBD | TBD |
| MACN-Or-ct 8947 | Uruguay | 26/03/2002 | -32.14 | -53.73 | TBD | TBD |
| MACN-Or-ct 9051 | Central Argentina | 11/12/15 | -32.59 | -69.35 | TBD | TBD |
| MACN-Or-ct 9055 | Central Argentina | 8/1/16 | -32.59 | -69.35 | TBD | TBD |
| MACN-Or-ct 9059 | Central Argentina | 21/01/2016 | -32.59 | -69.35 | TBD | TBD |
| MACN-Or-ct 9063 | Central Argentina | 1/11/16 | -32.59 | -69.35 | TBD | TBD |
| MACN-Or-ct 9067 | Central Argentina | 11/12/16 | -32.59 | -69.35 | TBD | TBD |
| MACN-Or-ct 9071 | Central Argentina | 24/01/2017 | -32.59 | -69.35 | TBD | TBD |
| MACN-Or-ct 9075 | Central Argentina | 17/12/2016 | -32.59 | -69.35 | TBD | TBD |
| MACN-Or-ct 9079 | Central Argentina | 23/12/2016 | -32.59 | -69.35 | TBD | TBD |
| MACN-Or-ct 9083 | Central Argentina | 21/01/2017 | -32.59 | -69.35 | TBD | TBD |
| MACN-Or-ct 9087 | Central Argentina | 28/01/2017 | -32.59 | -69.35 | TBD | TBD |
| MACN-Or-ct 9091 | Central Argentina | 29/01/2017 | -32.59 | -69.35 | TBD | TBD |
| MACN-Or-ct 9095 | Central Argentina | 4/2/17 | -32.59 | -69.35 | TBD | TBD |

| | | | | | | |
|---|---|---|---|---|---|---|
| MACN-Or-ct 9097 | Central Argentina | 2/12/15 | -32.59 | -69.35 | TBD | TBD |
| MACN-Or-ct 9099 | Central Argentina | 26/12/2015 | -32.59 | -69.35 | TBD | TBD |
| MACN-Or-ct 9103 | Northern Argentina | | -25.68 | -54.44 | TBD | TBD |
| MACN-Or-ct 9142 | Central Argentina | 7/11/17 | -34.55 | -59.44 | TBD | TBD |
| MACN-Or-ct 9144 | Central Argentina | 7/11/17 | -34.55 | -59.44 | TBD | TBD |
| QUMEL CW1 | MFI | 2009 | -52.2 | -59.15 | JQ943377 | TBD |
| QUMEL CW10 | MFI | 2009 | -52.2 | -59.15 | JQ943378 | TBD |
| QUMEL CW11 | MFI | 2009 | -52.2 | -59.15 | HM905961 | TBD |
| QUMEL CW2 | MFI | 2009 | -52.2 | -59.15 | HM905954 | TBD |
| QUMEL CW3 | MFI | 2009 | -52.2 | -59.15 | JQ943379 | TBD |
| QUMEL CW4 | MFI | 2009 | -52.2 | -59.15 | HM905955 | TBD |
| QUMEL CW5 | MFI | 2009 | -52.2 | -59.15 | HM905956 | TBD |
| QUMEL CW6 | MFI | 2009 | -52.2 | -59.15 | HM905957 | TBD |
| QUMEL CW7 | MFI | 2009 | -52.2 | -59.15 | HM905958 | TBD |
| QUMEL CW8 | MFI | 2009 | -52.2 | -59.15 | HM905959 | TBD |
| QUMEL CW9 | MFI | 2009 | -52.2 | -59.15 | HM905960 | TBD |
| UWBM 70276 | Northern Argentina | 2006 | -25.57 | -54.34 | HM396317 | TBD |
| UWBM 70288 | Northern Argentina | 1995 | -27.36 | -55.9 | HM396318 | TBD |

**Table S2. Genetic information for Troglodytes musculus/cobbi samples across southern South America.** Samples in bold indicate individuals with both mtDNA and RADseq data, the individual marked with "*" has only RADseq data and the remaining individuals in regular font only have mtDNA data (COI). The table includes the population of origin and its corresponding genetic cluster based on population structure analyses of mitochondrial and genomic data. For both the full dataset and a subset of loci used for linkage disequilibrium (LD) analysis, the table reports the number of sites retained, mean sequencing depth and proportion of missing data.

| Sample ID | Population | mtDNA cluster | Nuclear Genomic Cluster | # Sites | Mean depth | Missing | # Sites LD | Mean depth LD | Missing LD |
|---|---|---|---|---|---|---|---|---|---|
| **MACN-Or-ct 1016** | Northern Argentina | Widespread | Widespread | 6569 | 28.5 | 0.01 | 3698 | 28.5 | 0.01 |
| **MACN-Or-ct 1460** | Northern Argentina | Widespread | Widespread | 6565 | 57.7 | 0.01 | 3700 | 58.2 | 0.01 |
| **MACN-Or-ct 1475** | Northern Argentina | Widespread | Widespread | 6540 | 40.8 | 0.02 | 3693 | 41.2 | 0.01 |
| **MACN-Or-ct 1631** | Northern Argentina | Widespread | Widespread | 6599 | 45.7 | 0.01 | 3713 | 46.3 | 0.01 |
| **MACN-Or-ct 1710** | Northern Argentina | Patagonia | Patagonia | 6590 | 47.2 | 0.01 | 3712 | 47.6 | 0.01 |
| **MACN-Or-ct 1860** | Northern Argentina | Widespread | Widespread | 6580 | 30.7 | 0.01 | 3713 | 30.9 | 0.01 |
| **MACN-Or-ct 1895** | Central Argentina | Widespread | Widespread | 6570 | 66 | 0.01 | 3700 | 66.7 | 0.01 |
| **MACN-Or-ct 1902** | Central Argentina | Widespread | Widespread | 6572 | 51 | 0.01 | 3698 | 51.6 | 0.01 |
| **MACN-Or-ct 1903** | Central Argentina | Widespread | Widespread | 6570 | 41.2 | 0.01 | 3701 | 41.3 | 0.01 |
| **MACN-Or-ct 2333** | Patagonia | Patagonia | Patagonia | 6337 | 47.8 | 0.05 | 3568 | 48.1 | 0.05 |
| **MACN-Or-ct 2412** | Northern Argentina | Widespread | Widespread | 6548 | 34.1 | 0.02 | 3686 | 34.2 | 0.02 |
| **MACN-Or-ct 2586** | Patagonia | Patagonia | Patagonia | 6453 | 42.7 | 0.03 | 3634 | 43 | 0.03 |
| **MACN-Or-ct 2599** | Patagonia | Patagonia | Patagonia | 6574 | 46.3 | 0.01 | 3704 | 46.6 | 0.01 |
| **MACN-Or-ct 2624** | Patagonia | Patagonia | Patagonia | 6542 | 33.4 | 0.02 | 3684 | 33.7 | 0.02 |
| **MACN-Or-ct 2666** | Patagonia | Patagonia | Patagonia | 6515 | 59 | 0.02 | 3661 | 59.2 | 0.02 |
| **MACN-Or-ct 2679** | Patagonia | Patagonia | Patagonia | 6546 | 35.5 | 0.02 | 3681 | 35.7 | 0.02 |
| **MACN-Or-ct 2680** | Patagonia | Patagonia | Patagonia | 6502 | 48.3 | 0.02 | 3662 | 49 | 0.02 |
| **MACN-Or-ct 2846** | Northern Argentina | Northern Argentina | Widespread | 6270 | 27.5 | 0.06 | 3516 | 27.7 | 0.06 |
| **MACN-Or-ct 286** | Patagonia | Patagonia | Patagonia | 6574 | 57.4 | 0.01 | 3710 | 58.5 | 0.01 |

| | | | | | | | | | |
|---|---|---|---|---|---|---|---|---|---|
| **MACN-Or-ct 2893** | Northern Argentina | Northern Argentina | Widespread | 6580 | 43.8 | 0.01 | 3703 | 44.2 | 0.01 |
| **MACN-Or-ct 3168** | Northern Argentina | Widespread | Widespread | 6585 | 68.2 | 0.01 | 3706 | 69.2 | 0.01 |
| **MACN-Or-ct 3196** | Northern Argentina | Northern Argentina | Widespread | 6589 | 41.6 | 0.01 | 3710 | 41.8 | 0.01 |
| **MACN-Or-ct 3335** | Central Argentina | Widespread | Widespread | 6602 | 46.6 | 0.01 | 3719 | 47 | 0.01 |
| **MACN-Or-ct 3546** | Northern Argentina | Widespread | Widespread | 6578 | 49.2 | 0.01 | 3698 | 49.7 | 0.01 |
| **MACN-Or-ct 3636** | Central Argentina | Widespread | Widespread | 6057 | 6.6 | 0.09 | 3409 | 6.7 | 0.09 |
| **MACN-Or-ct 3759** | Patagonia | Patagonia | Patagonia | 6598 | 75 | 0.01 | 3718 | 75.6 | 0.01 |
| **MACN-Or-ct 3884** | Lowland Bolivia | Widespread | Widespread | 6573 | 116.6 | 0.01 | 3704 | 117 | 0.01 |
| **MACN-Or-ct 4100** | Central Argentina | Widespread | Widespread | 6599 | 52 | 0.01 | 3717 | 52.7 | 0.01 |
| **MACN-Or-ct 4162** | Northern Argentina | Widespread | Widespread | 6591 | 46.1 | 0.01 | 3713 | 46.6 | 0.01 |
| **MACN-Or-ct 4179** | Northern Argentina | Widespread | Widespread | 6437 | 19.8 | 0.03 | 3620 | 19.9 | 0.03 |
| **MACN-Or-ct 4204** | Northern Argentina | Widespread | Widespread | 6563 | 37.4 | 0.01 | 3692 | 37.7 | 0.01 |
| **MACN-Or-ct 4248** | Central Argentina | Widespread | Widespread | 6588 | 44.9 | 0.01 | 3704 | 45.2 | 0.01 |
| **MACN-Or-ct 4271** | Central Argentina | Widespread | Widespread | 6595 | 41.4 | 0.01 | 3711 | 41.5 | 0.01 |
| **MACN-Or-ct 4344** | Central Argentina | Widespread | Widespread | 6588 | 74.8 | 0.01 | 3708 | 75.3 | 0.01 |
| **MACN-Or-ct 4345** | Central Argentina | Widespread | Widespread | 6613 | 67.6 | 0.01 | 3726 | 68.1 | 0.01 |
| **MACN-Or-ct 4355** | Central Argentina | Widespread | Widespread | 6582 | 46.3 | 0.01 | 3701 | 46.2 | 0.01 |
| **MACN-Or-ct 4516** | Northern Argentina | Widespread | Widespread | 6579 | 49.6 | 0.01 | 3701 | 49.8 | 0.01 |
| **MACN-Or-ct 4560** | Patagonia | Patagonia | Patagonia | 6585 | 45.4 | 0.01 | 3712 | 45.4 | 0.01 |
| **MACN-Or-ct 4561** | Patagonia | Patagonia | Patagonia | 6598 | 37.1 | 0.01 | 3715 | 37.3 | 0.01 |
| **MACN-Or-ct 4563** | Central Argentina | Widespread | Widespread | 6600 | 58.2 | 0.01 | 3714 | 58.5 | 0.01 |
| **MACN-Or-ct 4567** | Central Argentina | Widespread | Widespread | 6556 | 24.6 | 0.01 | 3687 | 24.8 | 0.02 |
| **MACN-Or-ct 4579** | Central Argentina | Widespread | Widespread | 6572 | 35 | 0.01 | 3701 | 35.1 | 0.01 |
| **MACN-Or-ct 4663** | Lowland Bolivia | Widespread | Widespread | 6548 | 58.3 | 0.02 | 3686 | 58.8 | 0.02 |
| **MACN-Or-ct 4785** | Northern Argentina | Patagonia | Patagonia | 6576 | 88.8 | 0.01 | 3702 | 89.6 | 0.01 |
| **MACN-Or-ct 4793** | Northern Argentina | Patagonia | Patagonia | 6587 | 46.1 | 0.01 | 3701 | 46.5 | 0.01 |
| **MACN-Or-ct 4809** | Northern Argentina | Patagonia | Patagonia | 6542 | 44.9 | 0.02 | 3686 | 44.8 | 0.02 |
| **MACN-Or-ct 4811** | Northern Argentina | Widespread | Widespread | 6575 | 90.5 | 0.01 | 3699 | 91.1 | 0.01 |
| **MACN-Or-ct 5016** | Northern Argentina | Patagonia | Patagonia | 6574 | 39.5 | 0.01 | 3700 | 39.8 | 0.01 |

| | | | | | | | | | |
|---|---|---|---|---|---|---|---|---|---|
| **MACN-Or-ct 5063** | Northern Argentina | Widespread | Widespread | 6574 | 81.2 | 0.01 | 3699 | 81.6 | 0.01 |
| **MACN-Or-ct 5064** | Northern Argentina | Widespread | Widespread | 6593 | 70 | 0.01 | 3719 | 70 | 0.01 |
| **MACN-Or-ct 5077** | Northern Argentina | Northern Argentina | Widespread | 6585 | 76.4 | 0.01 | 3705 | 76.2 | 0.01 |
| **MACN-Or-ct 5101** | Central Argentina | Widespread | Widespread | 6598 | 61.7 | 0.01 | 3708 | 61.7 | 0.01 |
| **MACN-Or-ct 5125** | Central Argentina | Widespread | Widespread | 6614 | 74.4 | 0.01 | 3724 | 74.9 | 0.01 |
| **MACN-Or-ct 5326** | Central Argentina | Patagonia | Patagonia | 6608 | 72.9 | 0.01 | 3716 | 73.2 | 0.01 |
| **MACN-Or-ct 5327** | Central Argentina | Widespread | Widespread | 6592 | 89.8 | 0.01 | 3717 | 90.5 | 0.01 |
| **MACN-Or-ct 5336** | Central Argentina | Widespread | Widespread | 6496 | 28 | 0.02 | 3656 | 28.1 | 0.02 |
| **MACN-Or-ct 5384** | Northern Argentina | Widespread | Widespread | 6525 | 113 | 0.02 | 3684 | 114.9 | 0.02 |
| **MACN-Or-ct 55** | Patagonia | Widespread | Widespread | 6549 | 24.9 | 0.02 | 3692 | 25.4 | 0.01 |
| **MACN-Or-ct 5669** | Northern Argentina | Widespread | Widespread | 6532 | 23.9 | 0.02 | 3675 | 24.2 | 0.02 |
| **MACN-Or-ct 5684** | Northern Argentina | Widespread | Widespread | 6588 | 61.3 | 0.01 | 3704 | 61.5 | 0.01 |
| **MACN-Or-ct 5896** | Lowland Bolivia | Widespread | Widespread | 6290 | 36 | 0.05 | 3547 | 36.2 | 0.05 |
| **MACN-Or-ct 599** | Patagonia | Patagonia | Patagonia | 6567 | 37.1 | 0.01 | 3697 | 37.6 | 0.01 |
| **MACN-Or-ct 6112** | Central Argentina | Patagonia | Patagonia | 6598 | 30.5 | 0.01 | 3715 | 30.9 | 0.01 |
| **MACN-Or-ct 6126** | Central Argentina | Patagonia | Patagonia | 6511 | 19.2 | 0.02 | 3661 | 19.5 | 0.02 |
| **MACN-Or-ct 6178** | Northern Argentina | Widespread | Widespread | 6561 | 46.3 | 0.01 | 3692 | 46.8 | 0.01 |
| **MACN-Or-ct 6279** | Lowland Bolivia | Widespread | Widespread | 6458 | 28.1 | 0.03 | 3636 | 28.1 | 0.03 |
| **MACN-Or-ct 6386** | Patagonia | Patagonia | Patagonia | 6606 | 44.6 | 0.01 | 3717 | 44.6 | 0.01 |
| **MACN-Or-ct 6701** | Highland Bolivia | Highland Bolivia | Highland Bolivia | 6519 | 40.3 | 0.02 | 3671 | 40.5 | 0.02 |
| **MACN-Or-ct 6702** | Highland Bolivia | Highland Bolivia | Highland Bolivia | 6548 | 70.9 | 0.02 | 3684 | 71 | 0.02 |
| **MACN-Or-ct 6720** | Highland Bolivia | Highland Bolivia | Highland Bolivia | 6526 | 38 | 0.02 | 3673 | 38.1 | 0.02 |
| **MACN-Or-ct 6721** | Highland Bolivia | Widespread | Highland Bolivia | 6501 | 61.1 | 0.02 | 3662 | 61.8 | 0.02 |
| **MACN-Or-ct 6722** | Highland Bolivia | Highland Bolivia | Highland Bolivia | 6485 | 40 | 0.02 | 3655 | 40.3 | 0.02 |
| **MACN-Or-ct 6928** | Northern Argentina | Widespread | Widespread | 6585 | 85.4 | 0.01 | 3709 | 85.7 | 0.01 |
| **MACN-Or-ct 6937** | Northern Argentina | Widespread | Widespread | 6230 | 27.9 | 0.06 | 3515 | 27.9 | 0.06 |
| **MACN-Or-ct 695** | Patagonia | Patagonia | Patagonia | 6580 | 44 | 0.01 | 3703 | 44.3 | 0.01 |
| **MACN-Or-ct 7022** | Central Argentina | Widespread | Widespread | 6575 | 50 | 0.01 | 3697 | 50.1 | 0.01 |
| **MACN-Or-ct 7065** | Central Argentina | Patagonia | Widespread | 6555 | 43.3 | 0.01 | 3684 | 43.5 | 0.02 |

| | | | | | | | | | |
|---|---|---|---|---|---|---|---|---|---|
| **MACN-Or-ct 7528** | Northern Argentina | Widespread | Widespread | 6085 | 19.1 | 0.08 | 3425 | 19.4 | 0.09 |
| **MACN-Or-ct 7554** | Central Argentina | Patagonia | Patagonia | 6356 | 44.3 | 0.04 | 3569 | 44.7 | 0.05 |
| **MACN-Or-ct 7577** | Central Argentina | Widespread | Widespread | 6548 | 27.9 | 0.02 | 3685 | 28.1 | 0.02 |
| **MACN-Or-ct 7629** | Central Argentina | Widespread | Widespread | 6548 | 29.9 | 0.02 | 3686 | 30.3 | 0.02 |
| **MACN-Or-ct 7672** | Patagonia | Patagonia | Patagonia | 6514 | 48.2 | 0.02 | 3679 | 47.5 | 0.02 |
| **MACN-Or-ct 7702** | Central Argentina | Widespread | Widespread | 6542 | 25.9 | 0.02 | 3683 | 26.2 | 0.02 |
| **MACN-Or-ct 7736** | Northern Argentina | Widespread | Widespread | 6502 | 35 | 0.02 | 3668 | 34.6 | 0.02 |
| **MACN-Or-ct 7849** | Central Argentina | Widespread | Widespread | 6600 | 74.4 | 0.01 | 3712 | 75.1 | 0.01 |
| **MACN-Or-ct 7872** | Central Argentina | Widespread | Widespread | 6563 | 50.1 | 0.01 | 3700 | 50.5 | 0.01 |
| **MACN-Or-ct 8269** | Northern Argentina | Widespread | Widespread | 6465 | 27.7 | 0.03 | 3634 | 27.6 | 0.03 |
| MACN-Or-ct 8277* | Central Argentina | Missing | Widespread | 6534 | 44.9 | 0.02 | 3681 | 44.9 | 0.02 |
| **MACN-Or-ct 8674** | Uruguay | Widespread | Widespread | 5600 | 4.1 | 0.16 | 3121 | 4.1 | 0.17 |
| **MACN-Or-ct 8772** | Uruguay | Widespread | Widespread | 6361 | 10.4 | 0.04 | 3593 | 10.4 | 0.04 |
| **MACN-Or-ct 8916** | Uruguay | Widespread | Widespread | 6343 | 10.3 | 0.05 | 3577 | 10.4 | 0.04 |
| **MACN-Or-ct 8947** | Uruguay | Widespread | Widespread | 5630 | 5.8 | 0.15 | 3167 | 5.8 | 0.15 |
| **MACN-Or-ct 9051** | Central Argentina | Patagonia | Mendoza (Admix.) | 6518 | 35.2 | 0.02 | 3666 | 35.2 | 0.02 |
| **MACN-Or-ct 9055** | Central Argentina | Patagonia | Mendoza (Admix.) | 6421 | 30.1 | 0.03 | 3618 | 30.1 | 0.03 |
| **MACN-Or-ct 9059** | Central Argentina | Widespread | Mendoza (Admix.) | 6532 | 53.5 | 0.02 | 3691 | 53.5 | 0.01 |
| **MACN-Or-ct 9063** | Central Argentina | Patagonia | Mendoza (Admix.) | 6508 | 43.6 | 0.02 | 3667 | 43.6 | 0.02 |
| **MACN-Or-ct 9067** | Central Argentina | Patagonia | Mendoza (Admix.) | 6563 | 55 | 0.01 | 3705 | 54.6 | 0.01 |
| **MACN-Or-ct 9071** | Central Argentina | Patagonia | Mendoza (Admix.) | 6556 | 40.2 | 0.01 | 3692 | 40.3 | 0.01 |
| **MACN-Or-ct 9075** | Central Argentina | Patagonia | Mendoza (Admix.) | 6521 | 43.5 | 0.02 | 3662 | 43.6 | 0.02 |
| **MACN-Or-ct 9079** | Central Argentina | Widespread | Mendoza (Admix.) | 6527 | 49.1 | 0.02 | 3680 | 49.2 | 0.02 |
| **MACN-Or-ct 9083** | Central Argentina | Patagonia | Mendoza (Admix.) | 6591 | 48.4 | 0.01 | 3714 | 48.4 | 0.01 |
| **MACN-Or-ct 9087** | Central Argentina | Widespread | Mendoza (Admix.) | 6527 | 41.1 | 0.02 | 3673 | 40.9 | 0.02 |
| **MACN-Or-ct 9091** | Central Argentina | Widespread | Mendoza (Admix.) | 6545 | 42 | 0.02 | 3687 | 41.9 | 0.02 |
| **MACN-Or-ct 9095** | Central Argentina | Widespread | Mendoza (Admix.) | 6575 | 44.4 | 0.01 | 3698 | 44.2 | 0.01 |
| **MACN-Or-ct 9097** | Central Argentina | Widespread | Mendoza (Admix.) | 6571 | 50.7 | 0.01 | 3690 | 50.7 | 0.01 |
| **MACN-Or-ct 9099** | Central Argentina | Patagonia | Mendoza (Admix.) | 6407 | 28 | 0.04 | 3610 | 27.9 | 0.04 |

| | | | | | | | | | |
|---|---|---|---|---|---|---|---|---|---|
| MACN-Or-ct 9103 | Northern Argentina | Northern Argentina | Missing | - | - | - | - | - | - |
| **MACN-Or-ct 9142** | Central Argentina | Widespread | Widespread | 6570 | 24.1 | 0.01 | 3698 | 24.2 | 0.01 |
| **MACN-Or-ct 9144** | Central Argentina | Widespread | Widespread | 6524 | 37.6 | 0.02 | 3674 | 37.2 | 0.02 |
| **QUMEL CW1** | MFI | MFI | MFI | 5636 | 9.4 | 0.15 | 3161 | 9.3 | 0.16 |
| **QUMEL CW10** | MFI | MFI | MFI | 5580 | 7.7 | 0.16 | 3136 | 7.4 | 0.16 |
| **QUMEL CW11** | MFI | MFI | MFI | 6331 | 59.9 | 0.05 | 3573 | 59 | 0.05 |
| QUMEL CW2 | MFI | MFI | Missing | - | - | - | - | - | - |
| **QUMEL CW3** | MFI | MFI | MFI | 6161 | 24.8 | 0.07 | 3482 | 24.7 | 0.07 |
| **QUMEL CW4** | MFI | MFI | MFI | 6281 | 27.5 | 0.06 | 3538 | 27.1 | 0.06 |
| **QUMEL CW5** | MFI | MFI | MFI | 5493 | 26.4 | 0.17 | 3083 | 26 | 0.18 |
| QUMEL CW6 | MFI | MFI | Missing | - | - | - | - | - | - |
| **QUMEL CW7** | MFI | MFI | MFI | 6290 | 36.4 | 0.05 | 3536 | 36.3 | 0.06 |
| QUMEL CW8 | MFI | MFI | Missing | - | - | - | - | - | - |
| **QUMEL CW9** | MFI | MFI | MFI | 6240 | 23.6 | 0.06 | 3501 | 23.4 | 0.07 |
| UWBM 70276 | Northern Argentina | Northern Argentina | Missing | - | - | - | - | - | - |
| UWBM 70288 | Northern Argentina | Widespread | Missing | - | - | - | - | - | - |

31

**Table S3. G-PhoCS posterior probability estimates of demographic parameters among pairwise Southern House Wren and Cobb's Wren populations.** Estimates for effective population sizes (*Ne*), divergence times, and bidirectional migration rates among population pairs. For each pairwise comparison, the table reports the posterior probability median estimates followed by 95% credible intervals in parentheses. Effective population sizes (*Ne*) are shown in thousands (k), divergence times are given in thousands of years ago (kya), and migration rates (m) represent the proportion of individuals per generation moving between populations. The Effective Sample Size (ESS) for each parameter is indicated in square brackets. Scaled results for *Ne* and divergence times are reported using two mutation rates: $4.6 \times 10^{-9}$ substitutions per site per generation, estimated for flycatchers (Smeds et al. 2016), and $1.2 \times 10^{-9}$ substitutions per site per generation, estimated for Galliformes (Ellegren 2007).

| Pairwise comp. | mutation rate | Ne Pop1 (k) [ESS] | Ne Pop2 (k) [ESS] | Ne Ancestor (k) [ESS] | Divergence Time (kya) [ESS] | Migration m1→2 [ESS] | Migration m2→1 [ESS] |
|---|---|---|---|---|---|---|---|
| MFI-HBOL | $1.2 \times 10^{-9}$ | 27.6 (7.3–60.9) [1005] | 742.1 (579.3–1005.6) [4380] | 826.0 (556.0–2629.8) [761] | 1069.8 (282.7–6704.6) [188] | 0.09 (0–0.21) [971] | 0.02 (0.004–0.05) [713] |
| | $4.6 \times 10^{-9}$ | 7.2 (1.9–15.9) [1005] | 193.6 (151.1–262.3) [4380] | 215.5 (145.1–686.0) [761] | 279.1 (73.7–1749.0) [188] | | |
| MFI-PAT | $1.2 \times 10^{-9}$ | 17.1 (3.4–41.2) [802] | 1219.7 (665.3–2092.6) [1682] | 587.4 (376.1–909.6) [1328] | 717.0 (112.1–3218.5) [165] | 0.11 (0–0.50) [313] | 0.03 (0.004–0.09) [468] |
| | $4.6 \times 10^{-9}$ | 4.5 (0.9–10.8) [802] | 318.2 (173.5–545.9) [1682] | 153.2 (98.1–237.3) [1328] | 187.0 (29.2–839.6) [165] | | |
| MFI-WIDE | $1.2 \times 10^{-9}$ | 19.8 (4.5–42.7) [625] | 2741.8 (1915.9–3760.8) [3206] | 588.1 (467.3–712.8) [3294] | 875.2 (553.9–1240.2) [624] | 0.50 (0.10–1.02) [428] | 0.03 (0.005–0.07) [143] |
| | $4.6 \times 10^{-9}$ | 5.1 (1.2–11.1) [625] | 715.3 (499.8–981.1) [3206] | 153.4 (121.9–185.9) [3294] | 228.3 (144.5–323.5) [624] | | |

| | | | | | | | |
|---|---|---|---|---|---|---|---|
| PAT-HBOL | $1.2 \times 10^{-9}$ | 1233.6 (864.0–1702.6) [7736] | 907.5 (672.0–1183.0) [10660] | 652.5 (533.1–779.1) [6089] | 550.8 (427.7–673.9) [5676] | 0 (0–0) [605906] | 0 (0–0) [605539] |
| | $4.6 \times 10^{-9}$ | 321.8 (225.4–444.1) [7736] | 236.8 (175.3–308.6) [10660] | 170.2 (139.1–203.2) [6089] | 143.7 (111.6–175.8) [5676] | | |
| PAT-WIDE | $1.2 \times 10^{-9}$ | 602.9 (228.3–1124.5) [1192] | 3139.7 (2003.2–4494.0) [3463] | 561.6 (400.7–717.6) [2054] | 638.3 (378.4–1016.7) [433] | 0 (0–0) [36959] | 1.6 (0.3–3.7) [172] |
| | $4.6 \times 10^{-9}$ | 157.3 (59.6–293.3) [1192] | 819.0 (522.6–1172.4) [3463] | 146.5 (104.5–187.2) [2054] | 166.5 (98.7–265.2) [433] | | |
| WIDE-HBOL | $1.2 \times 10^{-9}$ | 2862.7 (1964.2–3990.3) [5502] | 717.0 (442.1–982.3) [2945] | 635.3 (504.9–770.3) [3648] | 655.6 (464.2–884.2) [1093] | 0.30 (0.01–0.81) [105] | 0.5 (0.05–1.2) [152] |
| | $4.6 \times 10^{-9}$ | 746.8 (512.4–1040.9) [5502] | 187.0 (115.3–256.3) [2945] | 165.7 (131.7–201.0) [3648] | 171.0 (121.1–230.7) [1093] | | |

### *Supplementary Figures*

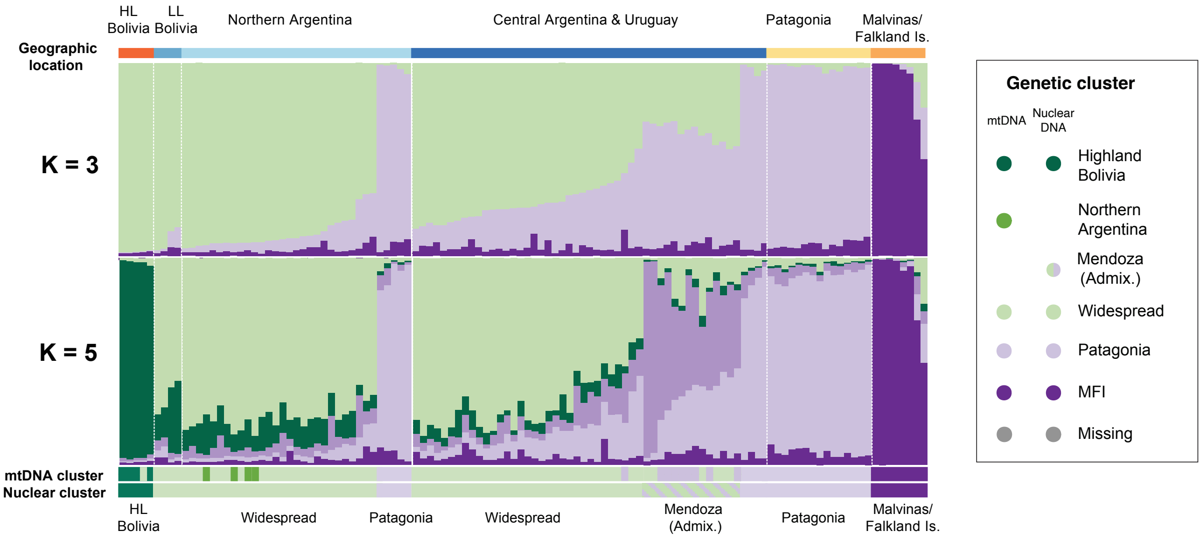

**Figure S1. Genetic structure of populations of the Southern and Cobb's Wrens across southern South America.** STRUCTURE analysis showing ancestry proportions for individuals assuming different numbers of genetic clusters (K = 3 and 5), with individuals ordered by geographic location: Highland Bolivia (HL), Lowland Bolivia (LL), Northern Argentina, Central Argentina & Uruguay, Patagonia, and the Malvinas/Falkland Islands. The bar in the bottom shows the genetic clustering according to fineRADstructure (see Figure 1E).

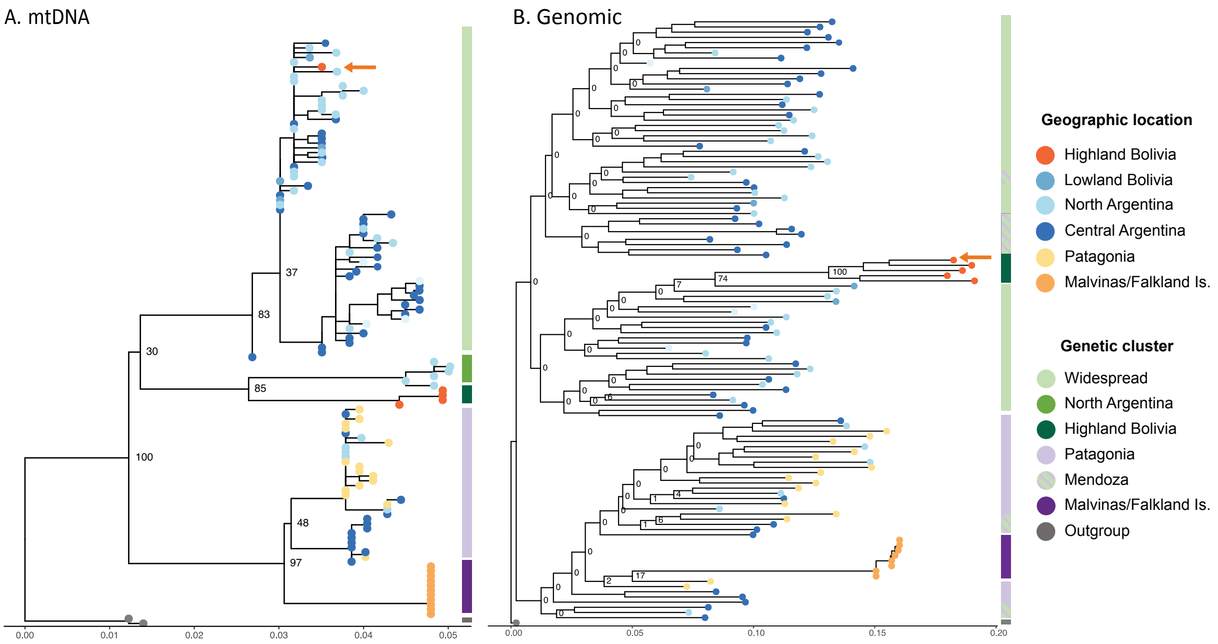

**Figure S2. Phylogenetic relationship among individuals based on 121 mtDNA COI (cytochrome oxidase I)** sequences and two outgroup individuals **(A)** and genome-wide SNP data of 116 individuals and one outgroup **(B)**, inferred using RAxML-NG. Tip colors indicate geographic origin, and the vertical color bars to the right of each tree denote genetic clusters. The x-axis represents the number of substitutions per site, reflecting genetic divergence among haplotypes. The orange arrow highlights an example of mito-nuclear discordance: one individual from the Highland Bolivia group clusters with Widespread individuals in the mtDNA tree.

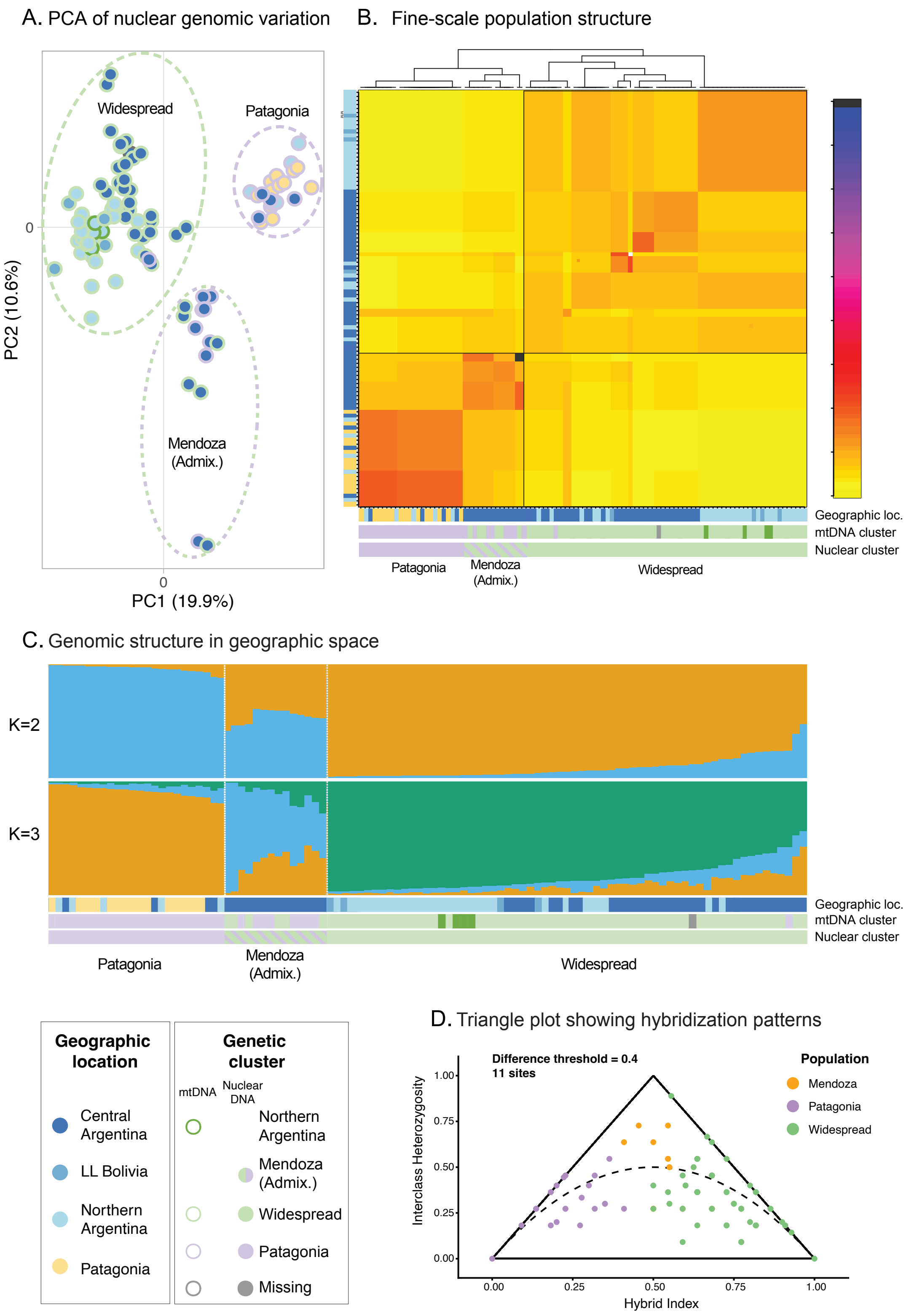
A. PCA of nuclear genomic variation
Widespread
Patagonia
Mendoza (Admix.)
PC2 (10.6%)
PC1 (19.9%)
0
0
B. Fine-scale population structure
Geographic loc.
mtDNA cluster
Nuclear cluster
Patagonia
Mendoza (Admix.)
Widespread
C. Genomic structure in geographic space
K=2
K=3
Geographic loc.
mtDNA cluster
Nuclear cluster
Patagonia
Mendoza (Admix.)
Widespread
Geographic location
Central Argentina
LL Bolivia
Northern Argentina
Patagonia
Genetic cluster
mtDNA
Nuclear DNA
Northern Argentina
Mendoza (Admix.)
Widespread
Patagonia
Missing
D. Triangle plot showing hybridization patterns
Difference threshold = 0.4
11 sites
Population
Mendoza
Patagonia
Widespread
Interclass Heterozygosity
Hybrid Index
1.00
0.75
0.50
0.25
0.00
0.00
0.25
0.50
0.75
1.00

**Figure S3. Genomic variation and evidence of admixture in the Mendoza population.** Principal component analysis (PCA) of nuclear genomic variation showing three main clusters: Widespread, Patagonia, and Mendoza (admixed). Circles are colored by geographic location, borders are colored by mtDNA genetic cluster, and dashed ellipses encompass individuals belonging to the same nuclear genetic cluster. **B)** fineRADstructure analysis showing the pairwise co-ancestry matrix with hierarchical clustering (top) and population assignments by geography (left and bottom bar) and mtDNA and genomic cluster (bottom bars).
**C)** STRUCTURE analysis showing ancestry proportions for individuals assuming two and three genetic clusters ($K = 2$ and $K = 3$), with individuals grouped by nuclear genetic cluster. Geographic location and genetic clusters inferred from mtDNA and nuclear DNA are shown in bars below. **D)** Triangle plot showing hybrid indices and interclass heterozygosity for individuals from the three clusters. The plot was generated using an allele frequency difference threshold of 0.4 between Patagonia and Widespread. Dashed lines represent expectations under different hybridization scenarios.
